# D-Serine agonism of GluN1-GluN3 NMDA receptors regulates the activity of enteric neurons and coordinates gut motility

**DOI:** 10.1101/2023.04.19.537136

**Authors:** Nancy Osorio, Magalie Martineau, Marina Fortea, Céline Rouget, Virginie Penalba, Cindy J. Lee, Werend Boesmans, Malvyne Rolli-Derkinderen, Amit V. Patel, Grégoire Mondielli, Sandrine Conrod, Vivien Labat-Gest, Amandine Papin, Jumpei Sasabe, Jonathan V. Sweedler, Pieter Vanden Berghe, Patrick Delmas, Jean-Pierre Mothet

**Affiliations:** Laboratoire de Neurosciences Cognitives (LNC), Aix-Marseille-Université, CNRS, UMR 7291, Marseille, France; Neurocentre Magendie, INSERM UMR U862, Bordeaux, France; Laboratory for Enteric NeuroScience (LENS), Translational Research Center for Gastrointestinal Disorders (TARGID), University of Leuven, Leuven, Belgium; Urosphere SAS, Toulouse, France; Department of Chemistry and the Beckman Institute for Advanced Science and Technology, University of Illinois at Urbana-Champaign, Urbana, IL, USA; Nantes Université, CHU Nantes, INSERM, The Institute of Digestive Diseases (IMAD), Nantes, France; Centre de Recherche en Neurophysiologie et Neuroscience de Marseille, UMR 7286, CNRS, Université Aix-Marseille, Marseille, France; Department of Pharmacology, Keio University School of Medicine, Tokyo, Japan; Université Paris-Saclay, École Normale Supérieure Paris-Saclay, Centre National de la Recherche Scientifique, CentraleSupélec, LuMIn UMR9024, Gif-sur-Yvette 91190, France

**Keywords:** Intestine, serine racemase, glutamate, enteric neurons, NMDA receptors

## Abstract

The enteric nervous system (ENS) is a complex network of diverse molecularly defined classes of neurons embedded in the gastrointestinal wall and responsible for controlling the major functions of the gut. As in the central nervous system, the vast array of ENS neurons is interconnected by chemical synapses. Despite several studies reporting the expression of ionotropic glutamate receptors in the ENS, their roles in the gut remain elusive. Here, by using an array of immunohistochemistry, molecular profiling and functional assays, we uncover a new role for D-serine (D-Ser) and non-conventional GluN1-GluN3 N-methyl D-aspartate receptors (NMDARs) in regulating ENS functions. We demonstrate that D-Ser is produced by serine racemase (SR) expressed in enteric neurons. By using both *in situ* patch clamp recording and calcium imaging, we show that D-Ser alone acts as an excitatory neurotransmitter in the ENS independently of the conventional GluN1-GluN2 NMDARs. Instead, D-Ser directly gates the non-conventional GluN1-GluN3 NMDARs in enteric neurons from both mouse and guinea-pig. Pharmacological inhibition or potentiation of GluN1-GluN3 NMDARs had opposite effects on mouse colonic motor activities, while genetically driven loss of SR impairs gut transit and fluid content of pellet output. Our results demonstrate the existence of native GluN1-GluN3 NMDARs in enteric neurons and open new perspectives on the exploration of excitatory D-Ser receptors in gut function and diseases.

## Introduction

The gastrointestinal (GI) tract and its auxiliary organs is an inherently complex ecosystem that is essential for life as it is the primary site for digestion of ingested food^1^. Therefore, GI functions require finely tuned control of muscular activity, fluid and electrolyte secretion and blood flow to efficiently propel the alimentary bolus, break down macroscopic food particles, and efficiently extract nutrients. Such control is accomplished by the enteric nervous system (ENS), which is intrinsically distributed throughout the GI wall^1, 2^. Although the ENS is regulated by the autonomic nervous system, it can uniquely function independently of the central nervous system (CNS)^3^. The importance of the ENS is emphasized by the life-threatening consequences of certain ENS congenital neuropathies, including chronic intestinal pseudo-obstruction or Hirschsprung’s disease^4, 5^ but also by its increasingly evident implication in inflammatory bowel diseases^6^ and brain disorders^7^.

The ENS is composed of the outer myenteric plexus and the inner submucosal plexus^1, 8^. The myenteric plexus is mainly involved in the control of GI peristalsis, whereas the submucosal plexus controls mucosal functions such as electrolyte secretion, paracellular permeability, and intestinal epithelial cell proliferation^1, 8, 9^. These ganglia are composed of diverse classes of neurons and glial cells that share many phenotypical features with their CNS counterparts, forming interlaced and expansive networks along the GI tract with defined reciprocal signaling^10–12^. Enteric neurons and glia cells use plethora of cell-cell signaling molecules to support GI functions^1, 12^. Virtually every class of neurotransmitters, and their receptors/transporters found in the CNS, has also been detected in the ENS^1, 9^. Accordingly, pathogenic mechanisms that give rise to CNS disorders might also lead to ENS dysfunction^7^.

Several studies have reported that glutamate can serve as a neurotransmitter in the ENS^13–17^. The idea that glutamate and its receptors are key signaling components in the GI arises from the observation that glutamate evokes fast and slow depolarizing responses of enteric neurons^13^. Subsequently, morphologic studies demonstrated the presence of some glutamatergic receptors in enteric neurons^14, 16, 18^ and functional studies have described the importance of glutamate in the modulation of acetylcholine release in the guinea-pig ileum and colon^15, 19^, as well as in motor and secretory functions in the gut^18, 20^. Therefore, alterations of glutamatergic transmission have been proposed to contribute to the pathogenesis of GI disorders^21, 22^.

Despite the consensus in the literature about glutamate as being a signaling molecule in the ENS^16, 21^, the identity of glutamatergic receptors involved in enteric neuronal network communication remains poorly defined^23^. Although the expression of functional α-amino-3-hydroxy-5-methyl-4-isoxazolepropionic acid receptors (AMPARs) and metabotropic glutamate receptors (mGluRs) in the ENS^14, 18, 24–26^ is well established, evidence for a functional role of the NMDA receptors (NMDARs) in the GI tract come from pharmacological studies in pathological conditions, where expression of these receptors is upregulated upon inflammation^27–29^. Whether and how NMDARs could contribute to the physiology of ENS in the small and large intestine remains to be established^26^.

NMDARs are heterotetrameric ligand-gated ion channels that associate two obligatory GluN1 subunits with two identical or different GluN2 or GluN3 or a mixture of GluN2 and GluN3 subunits^30, 31^. The activation of GluN1-GluN2 containing NMDARs requires the binding of glutamate plus a co-agonist glycine or D-Ser^30–32^ whereas diheteromeric GluN1-GluN3 receptors are not gated by glutamate^30, 31^. Mounting evidence gathered at central synapses indicate that D-Ser which is synthesized from L-Ser by serine racemase (SR)^33^ would be preferred over glycine thus controlling many NMDAR-dependent functions including synaptic plasticity and cognitive performances^34–37^.

While significant progress has been made in the understanding of the functions of D-Ser in the CNS, the role of D-Ser in the ENS is still unknown. However, indirect observations suggest that D-Ser may be an important signaling molecule in the gut. Indeed, acute kidney injury (AKI) induces gut dysbiosis by altering the metabolism of D-Ser while microbiota-derived D-Ser offers protection against AKI^38^. Furthermore, D-Ser produced by yet unidentified enteric cells regulates sleep in flies^39^. Whether D-Ser can be produced in the mammalian gut by host cells and can regulate the activity of the ENS and GI functions has remained unexplored.

Here, using a combination of pharmacological and genetic approaches, along with *in situ* patch clamp recording and calcium imaging, we have investigated the physiological role of glutamatergic signaling in the ENS. We report evidence that enteric neuron excitability and GI motility are not regulated by conventional GluN1-GluN2 containing NMDARs. Strikingly, we demonstrate that D-Ser is synthesized by SR in enteric neurons and gates the non-canonical GluN1-GluN3 receptors to shape ENS network activity for normal GI motility and transit.

## Results

### D-Serine is present in the gut and is metabolized by enteric neurons

We first assessed whether the amino acids (AAs) D-Ser and L-Ser are present within the mouse intestine. AAs were quantified by CE–LIF in the lumen and the wall (lacking the mucosa and submucosa) of the medial parts of mouse ileum and colon. **Figure 1a,b** reveal the presence of a substantial amount of D- and L-Ser in both the lumen and the wall fractions. The D-Ser peak identity was validated by spiking with D-Ser standards (not shown) and confirmed *via* enzymatic degradation by *Pk*DAAO, as the enzyme shows high catalytic activity towards D-Ser (insets in **Fig. 1a**). Given the abundance of microbiota-derived D-Ser in the gut^38, 40^, we then compared the levels of D-Ser in WT and in serine racemase (SR) null-mutant (SR^-/-^) mice. A statistically significant decrease of ∼50% was observed in D-Ser content of both ileum and colon from SR^-/-^ mice compared with WT mice (**Figure 1b**). Although free D-AAs in the murine intestinal tract may be microbial products^41^, our observations suggest that an important proportion of free D-Ser in the gut originate from its local biosynthesis by the host intestinal cells.

**Figure 1.**
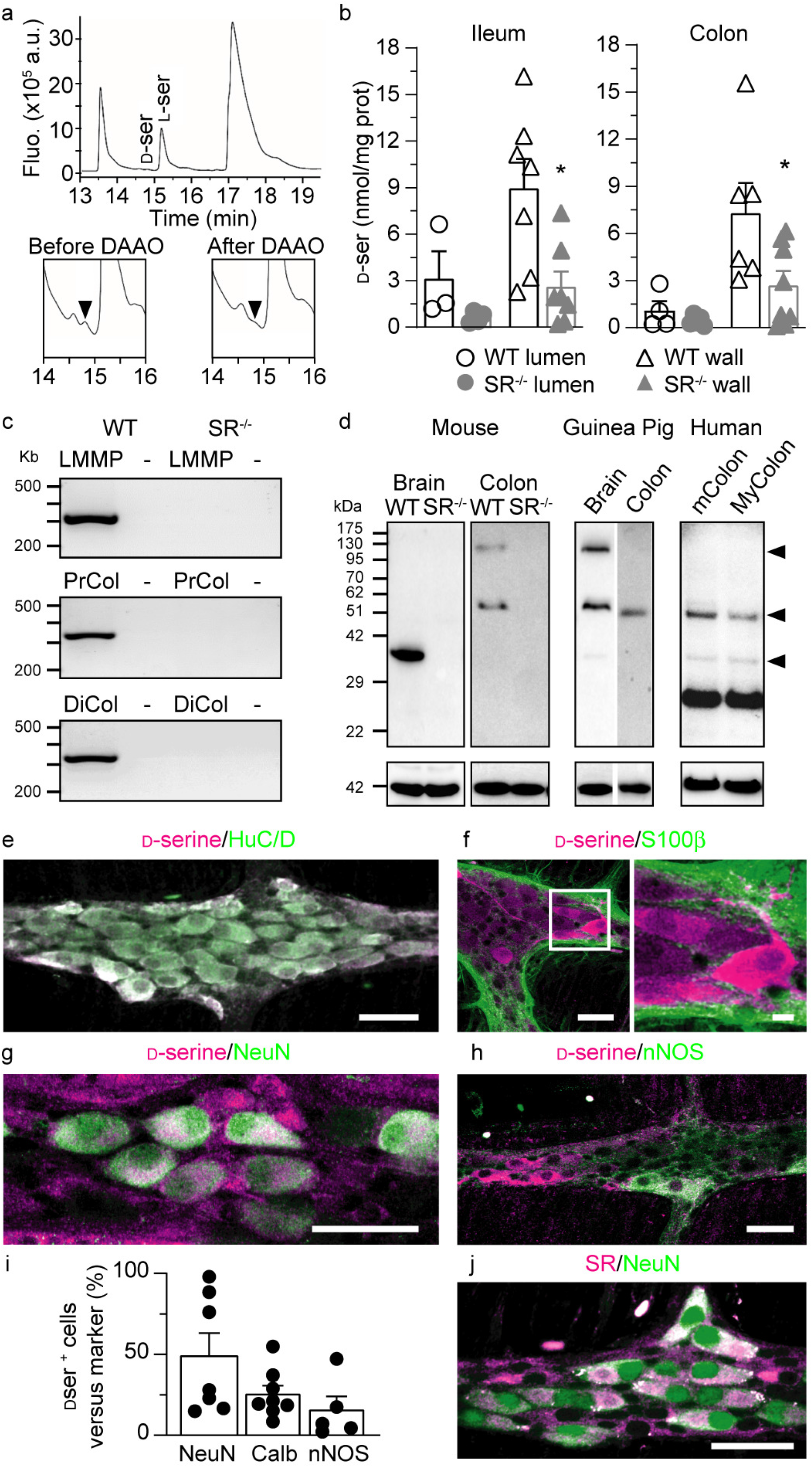
D-Serine and serine racemase are present in enteric neurons. **a**) Representative electropherogram of a NDA-derivatized gut sample showing the presence of two peaks corresponding to L-serine (L-Ser) and D-serine (D-Ser). The D-Ser peak identity was confirmed by enzymatic degradation by DAAO (lower panels). **b**) Histograms showing the content of D-Ser (mean ± s.e.m) found in the lumen and the wall of the medial parts of ileum (left panel) and colon (right panel) of WT (n = 4) and SR^-/-^ (n = 4) mice. *p<0.05 using one-way ANOVA followed by Tukey’s multiple comparison test. **c**) RT-PCR analysis showing the expression of SR transcripts in colonic LMMP (top) and proximal (PrCol) and distal (DiCol) colon sections of WT but not SR^-/-^ mice. (-) indicates no RT. **d**) Western blot analysis of the expression of SR in brain and colonic LMMP from WT and SR^-/-^ mice, brain and colon tissues from guinea pigs and the medial part of human colon with (mColon) and without (MyColon) the mucosa. Protein loading was normalized to actin (42 kDa). **e**) Immunostaining for D-Ser (magenta) and the neuronal pan marker HuC/D (green). N = 5 guinea-pigs, 5 preparations. Scale: 50 µm. **f**) Immunostaining for D-Ser (magenta) and the glial marker S100B (green). N = 4 guinea-pigs, 4 preparations. Scale: 50 µm left, 10 µm right. **g**) Immunostaining for D-Ser (magenta) and the marker of IPANs, NeuN (green). N = 7 guinea pigs, 7 preparations. Scale: 50 µm. **h**) Immunostaining for D-Ser (magenta) and the neuronal nitric oxide synthase (nNOS, green), a marker of inhibitory motoneurons. N=4 guinea-pigs, 5 preparations. Scale: 50 µm. **i**) Bar graph showing the mean percentage (+/-s.e.m) of D-Ser-positive myenteric neurons co-labelled with NeuN (7 guinea-pigs, 7 preparations), Calbindin (Calb, 8 guinea-pigs, 8 preparations) and nNOS (4 guinea-pigs, 5 preparations). **j**) Immunostaining for SR (magenta) and IPAN marker NeuN (green) (N = 5 guinea pigs, 5 preparations) Scale: 50 µm.

We therefore explored whether SR is present in the intestine to synthesize D-Ser. RT-PCR performed from mouse colon samples showed the presence of SR mRNA in the distal and proximal sections of the colon of WT mice, but not in tissues extracted from SR^-/-^ mice (**Figure 1c**). Interestingly, mRNA for SR was also detected in LMMP (Longitudinal Muscle and Myenteric Plexus) preparations of mouse colon, which lacks the circular muscle layer and the mucosa-submucosa but still comprises the myenteric neuronal network (**Figure 1c**). Western blot analysis confirmed that SR is present in the colon of WT mice as well as in guinea pigs and humans (**Figure 1d**). Although mouse brain SR was detected as a monomeric form at 37kDa, in all colon samples and guinea pig brain multiple bands of higher sizes were observed, which most likely corresponds to previously reported variants of SR^42^ or to unreducible multimers^43–47^. The absence of any signal from SR^-/-^ mice further demonstrated that our antibody was specific for SR and that the higher molecular weight bands were indeed caused by SR forms resistant to both denaturation and reductive treatment. The lower molecular weight bands in human samples may indicate the presence of proteolytic fragments of SR^48^.

To identify which cells in the ENS may synthesize D-Ser, we next performed immunostainings in the guinea pig ENS, the best molecularly characterized ENS model^8^. LMMP immunostainings for D-Ser revealed a high density of cell body labelling in the myenteric plexus of the ileum (**Figure 1e-j** **and Extended data Fig.1**). Double immunostainings with the pan neuronal marker HuC/D revealed the presence of D-Ser in neuronal cell bodies (**Figure 1e** and **Extended data Fig. 1a**). On the contrary, double labelling for the glial marker S100β or the glial fibrillary acidic protein (GFAP, data not shown) and D-Ser showed that D-Ser is not detected in glia cells (**Figure 1f** and **Extended data Fig. 1b**).

As the ENS comprises diverse molecularly and functionally defined classes of neurons, we next investigated the identity of the neurons containing D-Ser by comparing its distribution with known markers. Double immunostainings revealed that 49% of D-Ser^+^-neurons also express NeuN, an exclusive marker of Dogiel type II neurons, which are considered as intrinsic primary afferent neurons (IPANs) in the guinea-pig GI tract^49, 50^ (**Figure 1g,i** and **Extended data Fig. 1c**). Consistently calbindin, another preferential marker of IPANs, was also found to be expressed in a substantial proportion of D-Ser^+^ myenteric neurons (**Figure 1i**). Sixteen percent of D-Ser^+^ neurons were co-labelled with the neuronal nitric oxide synthase (nNOS), which is specific for inhibitory motoneurons and descending interneurons^8^ (**Figure 1h,i** and **Extended data Fig. 1d**).

Because the presence of D-Ser in neuronal cell bodies does not necessarily reflect its synthesis site, we stained LMMPs for SR. A very similar labelling profile was observed for SR and NeuN in the myenteric plexus, thus suggesting that IPANs are one of the main cells potentially producing D-Ser in the gut (**Figure 1j** and **Extended data Fig. 1e**). Finally, we asked whether the D-Amino Acid Oxidase (DAAO), which is degrading D-Ser^51^ and is highly expressed in the hindbrain, is also present in the mouse gut. We found DAAO to be present in the small intestine but not in other regions of the GI tract in agreement with previous data^41^ (**Extended data Fig. 2 & 3)**. Collectively these observations suggest that enteric neurons express SR and may produce D-Ser throughout the GI tract, pointing to a putative role of D-Ser in the regulation of ENS functions.

### Molecular profiling reveals the presence of multiple glutamatergic receptor subunits in the myenteric plexus

In addition to GluN1-GluN2 containing NMDAR, D-Ser can also bind to the ionotropic GluN1-GluN3 unconventional NMDARs^52, 53^ and be a ligand for the GluD1-D2 receptors^54–56^. To identify the putative receptor for D-Ser, we performed RT-PCR for the GluN1, GluN2A-D, GluN3A-B and GluD1-2 receptor subunits on LMMPs isolated from the colon of WT mice (**Extended data Fig. 4a**). Amongst the transcripts tested, only GluN1, GluN2D, GluN3A, GluA2, GluD1 and GluD2 transcripts were detected. The presence of receptor subunit proteins was further analyzed by immunoblotting with previously characterized antibodies (**Extended data Fig. 4b**). Consistent with RT-PCR data, immunoblots showed the presence of GluN1 and GluN2C/2D in the colon, whereas GluN2A/B were not detected in mouse LMMPs. As the antibody we used does not distinguish between GluN2C and GluN2D (**Extended data Fig. 4b, middle panel**), we conclude for the presence of GluN2D based on the results of RT-PCR, which show the presence of GluN2D but not GluN2C transcripts. Immunoblottings demonstrated the presence of GluN3A and GluN3B subunits in mouse colon (**Extended data Fig. 4b, middle panel**). Of note, the expression of the glutamatergic receptor subunits was unchanged in SR^-/-^ mice (**Extended data Fig. 4b)**.

Our data indicate that GluN1, GluN2D, GluN3A and GluN3B are detected in the mouse colon. The presence of these receptor subunits supports two main scenarios for a modulatory role of D-Ser. In the first scenario, D-Ser would bind to diheteromeric GluN1-GluN2D receptors or triheteromeric GluN1-GluN2D-GluN3 receptors, which both require glutamate for activation and are inhibited by the glutamate site competitive antagonist D-AP5^30, 31^. In the second scenario, D-Ser would bind to the unconventional diheteromeric GluN1-GluN3 NMDARs, which are not gated by glutamate or NMDA and not blocked by D-AP5^30, 31^.

### Enteric D-Ser does not act through post- or pre-synaptic GluN1-GluN2 containing NMDARs

We next performed *in situ* patch clamp recordings, both in voltage and current clamp, of myenteric neurons in guinea-pig LMMPs^57^ to first probe potential interaction of D-Ser with GluN1-GluN2 containing NMDARs.

In voltage clamp mode, we found that local application of glutamate evoked large post-synaptic inward currents in myenteric neurons (**Figure 2a,b**). However, these post-synaptic currents were evoked with relatively high concentrations of glutamate (20 mM) and were not blocked by the selective NMDAR antagonist D-AP5 (**Figure 2a**). In addition, the glutamate-induced current-voltage (I/V) relationship determined using slow voltage ramps lacked the typical negative slope of NMDAR currents due to voltage-dependent Mg^2+^ block (**Figure 2c**). The lack of functional GluN1-GluN2 NMDARs was further substantiated by the absence of responses to NMDA (100 µM-1 mM) in myenteric neurons that either responded to glutamate or kainate (**Figure 2d,e**). Additionally, application of NMDA in the continued presence of Tricine, which alleviates any blockade by endogenous Zn^2+^ of GluN2A-containing NMDARs^36^, did not evoke any currents at very positive potentials (**Figure 2e**).

**Figure 2.**
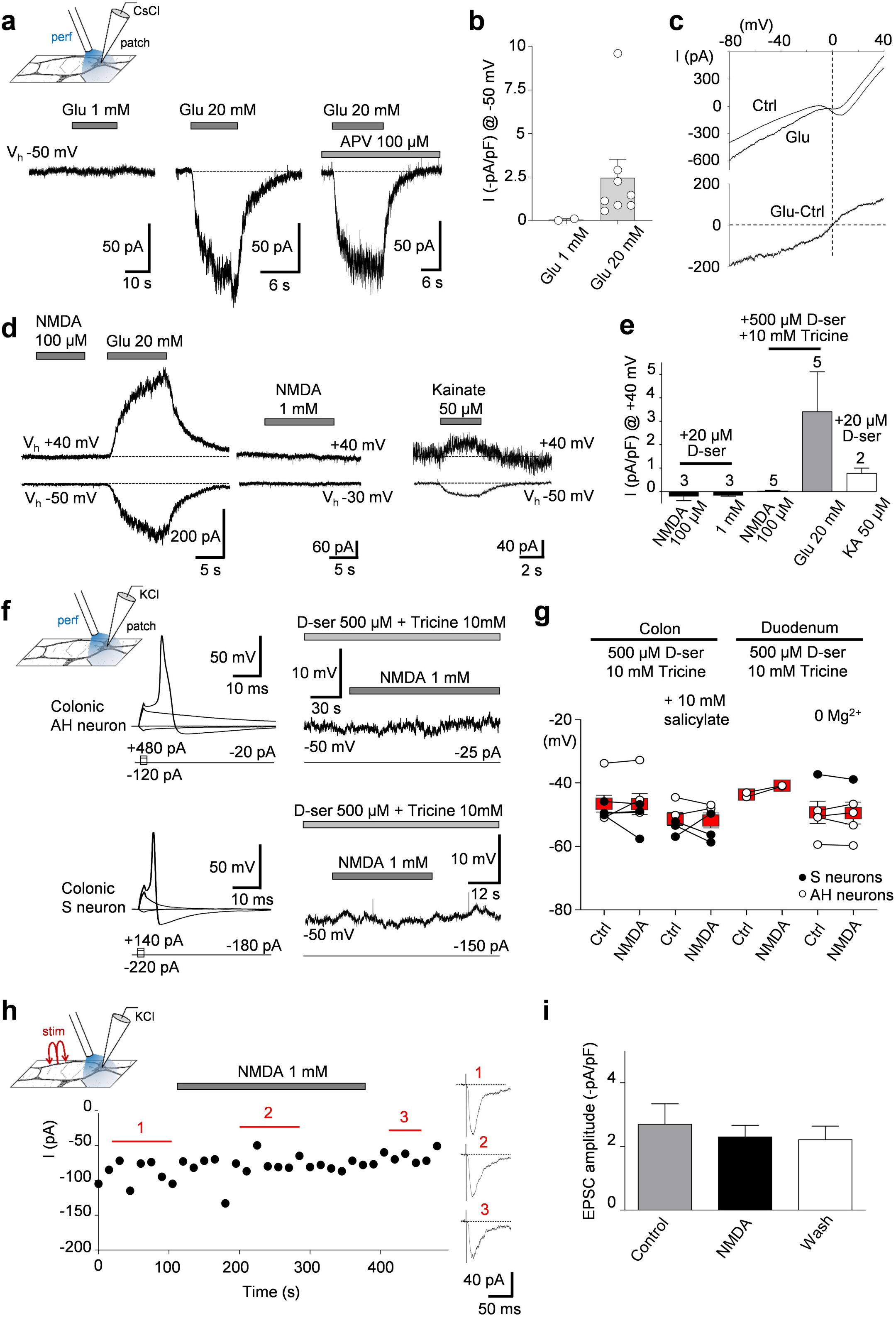
Lack of detectable functional NMDA receptors at both pre- and post-synaptic levels in myenteric neurons. **a**) Glutamate at 20 mM, but not at 1 mM, evoked inward currents that were insensitive to APV (D-AP5, 100 μM), a NMDA receptor antagonist. Recordings of *in situ* guinea-pig myenteric neurons were made in the voltage-clamp mode with a holding potential of -50 mV, in the presence of 500 µM D-Ser and 10 mM tricine in the bath medium. CsCl-based intrapipette solution. **b**) Quantification of 1 and 20 mM glutamate-evoked currents (I = 0.05 ± 0.01 pA/pF, n = 2 and I = 2.46 ± 1.06 pA/pF, n = 8, respectively). **c**) Upper traces: slow voltage-ramp induced *I-V* relationships determined before and during the application of 20 mM glutamate (Glu). Lower trace: difference current depicting the voltage-dependence of the glutamate-induced current. Note the absence of the N-shaped *I-V* relationship typical of NMDAR-induced current. Currents were evoked by a 2s-voltage ramp command from -80 to +40 mV (0.06 mV/ms) in the continued presence of 500 µM D-Ser, 10 mM tricine and 1 mM Mg^2+^ in the bath. **d**) Glutamate (20 mM) and kainate (50 µM) evoked currents at either negative or positive potentials, whereas NMDA, at 100 μM or 1 mM, failed to do so. Recordings were made in the voltage-clamp mode at the indicated holding potentials, in the presence of 500 µM D-Ser and 10 mM tricine in the bath medium. CsCl-based intrapipette solution. **e**) Pooled data showing Glutamate-, Kainate- and NMDA-induced currents determined at Vh = +40 mV. Recording conditions are indicated. From left to right: I_NMDA_ = -0.21 ± 0.19 pA/pF; I_NMDA_ = -0.17 ± 0.04 pA/pF; I_NMDA_ = 0.01 ± 0.04 pA/pF; I_Glu_ = 3.4 ± 1.7 pA/pF; I_KA_ = 0.78 ± 0.23 pA/pF. **f**) Recordings of membrane potential during 1 mM NMDA application in a colonic AH myenteric neuron (top trace) and a colonic S myenteric neuron (bottom panel). Current clamp recordings were made in the presence of 500 µM D-Ser and 10 mM tricine using K^+^-based pipette solution. AH- and S-neuronal subtypes were ascertained from their action potential waveform (left panels). Left: Evoked action potentials of the AH and S neurons to current injections of -2, 0, 16 and 20 pA/pF and -1.6, 0, 9.6 and 12.8 pA/pF respectively. **g**) Membrane potential measurements in S- and AH-type myenteric neurons from both guinea-pig colonic and duodenal myenteric neurons before and during superfusion of NMDA (1 mM) in the presence of 500 µM D-Ser and 10 mM tricine. KCl intrapipette solution. From left to right (mean +/-s.e.m, mV): Vm_ctrl_ = -46.6 ± 2.7, Vm_NMDA_ = -46.8 ± 3.3, n = 6; Vm_ctrl_ = -51.4 ± 2.1, Vm_NMDA_ = -51.9 ± 2.4, n = 5; Vm_ctrl_ = -43.7 ± 0.7, Vm_NMDA_ = -40.9 ± 0.1, n = 2; Vm_ctrl_ = -49.3 ± 3.5, Vm_NMDA_ = -49.6 ± 3.5, n = 5. **h**) Amplitude of presynaptically-evoked fast nicotinic EPSCs recorded in a GP myenteric neuron from duodenal LMMP before, during and after application of NMDA (1 mM). Holding potential -50 mV. Recordings made in the presence of 20 µM D-Ser and 10 mM tricine. KCl-based intrapipette solution. Single shock electrical stimulation was applied on nerve bundles every 15 s. Right insets: corresponding recordings (average of 4-7 sweeps) of EPSCs before (1), during (2) and after (3) application of NMDA. **i**) Mean amplitudes of electrically evoked EPSCs before (control), during (NMDA) and after (wash) application of NMDA (1 mM) in the presence of 20 µM D-Ser and 10 mM tricine. ICtrl = -2.7 ± 0.6 pA/pF; I_NMDA_ = -2.3 ± 0.4 pA/pF; I_wash_ = -2.2 ± 0.4 pA/pF (n = 3). Data were normalized to the cell membrane capacitance.

Current clamp recordings were used to test whether NMDA would alter the resting membrane potential in identified myenteric neuron populations. This recording configuration indeed allows for distinguishing IPANs from interneurons and motor neurons, also called S neurons^50^. IPANs are AH neurons, in which the action potential shows a shoulder, sustained by Ca^2+^ and Nav1.9 channel activation, and a slow afterhyperpolarizing (AH) potential, whereas S neurons have monophasic action potential repolarization and no slow afterhyperpolarization^58, 59^. Neither AH neurons, nor S neurons from the colon and the duodenum were found to be sensitive to NMDA exposure under a variety of conditions (D-Ser, Tricine, salicylate, free [Mg^2+^]o) that would promote NMDAR function (**Figure 2f,g**).

To examine whether GluN1-GluN2 NMDARs may be active at the presynaptic terminal, we coupled patch clamp recordings of postsynaptic myenteric neurons in LMMPs with electric stimulation of presynaptic nerve bundles. We applied single electric stimuli at low frequency (1 ms every 15 s), which evoked fast excitatory post-synaptic currents (fEPSCs) mediated by nicotinic receptors. Local application of NMDA failed to modulate the presynaptically-evoked fEPSCs (**Figure 2h,i**). Therefore, our electrophysiological recordings failed to detect functional GluN1-GluN2 containing NMDARs at both post- and pre-synaptic neurons, thus raising questions about the identity of the putative receptor for D-Ser.

### Acutely applied D-Ser raises intracellular calcium concentrations in colonic myenteric neurons

We next performed calcium imaging as readout for investigating the effects of D-Ser on a large population of myenteric neurons in LMMPs prepared from mouse colon. Using a local perfusion pipette, we exposed Fluo-4 loaded myenteric ganglia to 100 μM D-Ser for 20s (**Figure 3a1-a3**). This local D-Ser exposure evoked calcium responses in 30 % of myenteric neurons (range from 28 to 92%) with a mean increase of fluorescence change (ΔF/F0) of 20%. By comparison the fluorescence change induced by depolarization with high [K^+^]_o_ was about 10 fold higher (191%, not shown). The D-Ser-induced Ca^2+^ signals had a relatively slow onset, peaked after 10-20s and declined within 20s to resting values (**Figure 3a4**). The slow kinetics of the responses suggests that most D-Ser-induced Ca^2+^ signals were not associated with abrupt changes in membrane potential^12, 60^.

**Figure 3.**
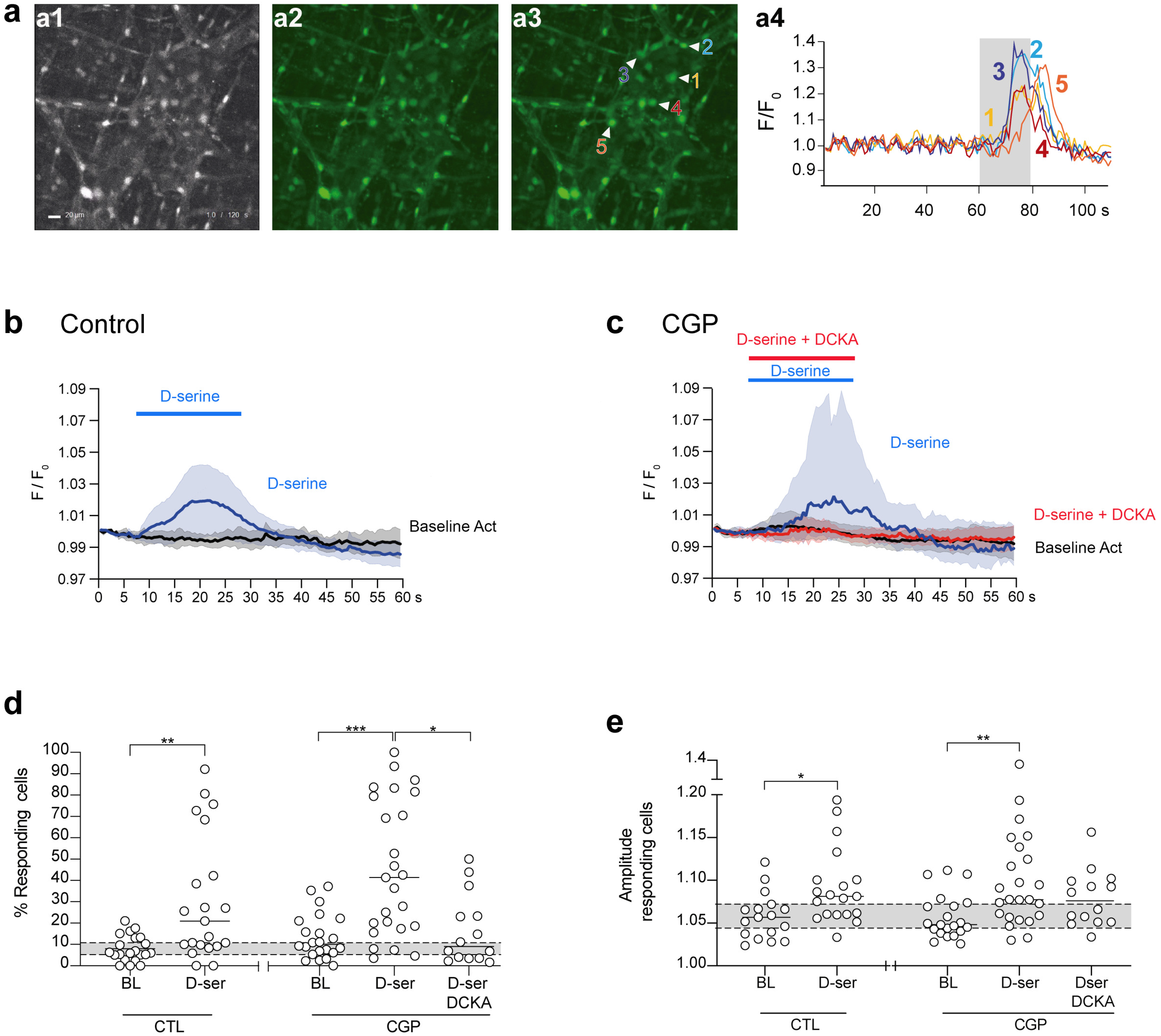
Acute application of D-Ser evokes Ca^2+^ responses in mouse myenteric neurons. **a**) Fluo-4 loaded colonic myenteric ganglia were exposed to D-Ser (100 μM, 20 s), which elicited a rise in intracellular [Ca^2+^]_i_ in some neurons. Scale = 20 µm. **a1**) Frame of a Fluo-4 movie in which at frame 60 a colonic ganglion is exposed to D-Ser for 20s. **a2-3**) Averages of 10 frames of the recording at rest (**a2**) and after exposure to D-Ser (**a3**). **a4**) Fluo-4 intensity traces of five individual actively responding neurons. The grey area indicates D-Ser application. **b**) Normalized fluorescence intensity of the average of D-Ser-induced [Ca^2+^]_i_ transients in control conditions (n = 16 neurons). (**c**) Averaged D-Ser-induced [Ca^2+^]_i_ transients (n = 23 neurons) in the continued presence of CGP-78608 (500 nM) (blue trace) and when CGP-78608 (500 nM) and DCKA (10 μM) were applied concomitantly. In **b** and **c**, graphs represent the mean ± 95% CI (as indicated by the shaded areas) of the normalized intensity over time. Colour-coded bars indicate the condition and the stimuli used, their beginning and duration. **d,e**) Percentage of responding cells (including spontaneous responses) per ganglion (**d**) and corresponding amplitude of the response per ganglion (**e**) to local perfusion of D-Ser in basal conditions (Krebs, BL) as well as after CGP-78608 or CGP-78608 +DCKA incubation. Data collected from 30-35 cells per ganglion and 6 mice for each condition. Each open circle refers to one ganglion. The grey bar represents the control baseline 95% CI. Horizontal lines indicate mean values. Unpaired t test or Mann-Whitney test were performed when comparing in groups. Significant *p* values are indicated in the graphs (* p ≤ 0.05, ** p ≤0.01, *** p ≤ 0.001).

To test the involvement of unconventional GluN1-GluN3 NMDARs we took advantage of the properties of the pharmacological compounds CGP-78608 and dichlorokynurenic acid (DCKA). While binding of D-Ser to the high-affinity GluN3 subunit triggers channel opening, D-Ser binding to the associated low-affinity GluN1 subunit has an opposite effect, causing auto-inhibition of the receptor^61, 62^. Thus, the CGP-78608 compound by preventing D-Ser binding on GluN1 can act as a potentiator of GluN1-GluN3 receptor^62^. Conversely, DCKA prevents D-Ser binding to both subunits and consequently blocks GluN1-GluN3 receptor activity. To avoid experimental bias caused by variability among preparations, new control experiments were included for every set of experiments to enable drug comparisons with control experiments performed at the same time and in the same animals. In this dataset local perfusion of D-Ser (100 µM) evoked transient increases in [Ca^2+^]_i_ in 30.9 ± 29.4% of Fluo-4 loaded neurons in each ganglion (**Figure 3b,d**). As in all datasets, this was significantly higher than the spontaneous and reverberating activity that occurred in the myenteric neuronal network (8.0 ± 6.1% neurons, p = 0.001) (**Figure 3d**). D-Ser-induced Ca^2+^ transients in myenteric neurons (Fi/F0: 1.09 ± 0.04) were significantly higher compared to baseline variations in amplitude (Fi/F0: 1.06 ± 0.03; p = 0.009) (**Figure 3b,d**). In the presence of CGP78608 (500 nM, preincubation for 15 min), the proportion of neurons responding to D-Ser increased to 45.5 ± 31.9%, which was significantly higher compared to the spontaneously active proportion (13.5 ± 10.7%, p = 0.0001) (**Figure 3c,d**). Also the [Ca^2+^]_i_ response amplitude was enhanced compared to baseline (p = 0.0019) (**Figure 3c,e**). Importantly, DCKA (10 μM) strongly reduced the proportion of D-Ser responding neurons in the continued presence of the CGP78608 from 45.5% to 16.6 ± 16.5% (p = 0.02) (**Figure 3c,d**). Equally, the increase of D-Ser-evoked Ca^2+^ transients in the presence of CGP78608 was prevented by the preincubation of DCKA (**Figure 3e**). Overall, these data support the notion that the GluN1-GluN3 receptors can be activated by D-Ser alone, generating Ca^2+^ rises independently of GluN2-containing receptors.

### D-Ser activates GluN1-GluN3 receptor-mediated slow excitatory currents in myenteric neurons

We used *in situ* patch-clamp recordings to unravel the exact mechanisms of GluN1-GluN3 receptor activation by D-Ser in mouse and guinea-pig myenteric neurons. The action of D-Ser was studied using the whole-cell voltage-clamp mode coupled to a local perfusion system positioned over the recorded neuron. In most mouse myenteric neurons tested (79%, 11/14 neurons), application of D-Ser alone (100 μM) caused fluctuations in the holding current that were difficult to distinguish from the intrinsic electrical fluctuations of the enteric neuronal network (**Figure 4a**). In a minority of cases (21%, 3/14), D-Ser elicited a small (-0.37 ± 0.03 pA/pF) slowly developing inward current, reminiscent of the typical glycine-activated GluN1-GluN3 NMDAR current in brain neurons^62^. Pretreatment with CGP78608 (500 nM) significantly increased the percentage of neurons that responded to D-Ser to 55% (11/20 neurons). In these cells, D-Ser-induced slow inward current reached a mean amplitude of -1.27 ± 0.18 pA/pF. Overall, the D-Ser inward current in CGP78608-treated neurons was potentiated 4 fold compared with non CGP78608-treated neurons (**Figure 4a,e**). D-Ser currents caused a mean depolarization of 6 mV in the receiving neuron’s membrane potential after CGP78608 treatment (**Figure 4d**). Pre-application of CGP78608 also increased the proportion of myenteric neurons exhibiting D-Ser-induced currents (from 13 to 42%) as well as the amplitude of these currents (from -0.15 ± 0.06 to -0.55 ± 0.07 pA/pF; p<0.01) in guinea-pig LMMPs (**Figure 4b,e**). Consistent with the involvement of GluN1-GluN3 receptors in D-Ser responses, DCKA fully prevented the occurrence of D-Ser-mediated inward currents in the continued presence of CGP78608 (**Figure 4c,e**).

**Figure 4.**
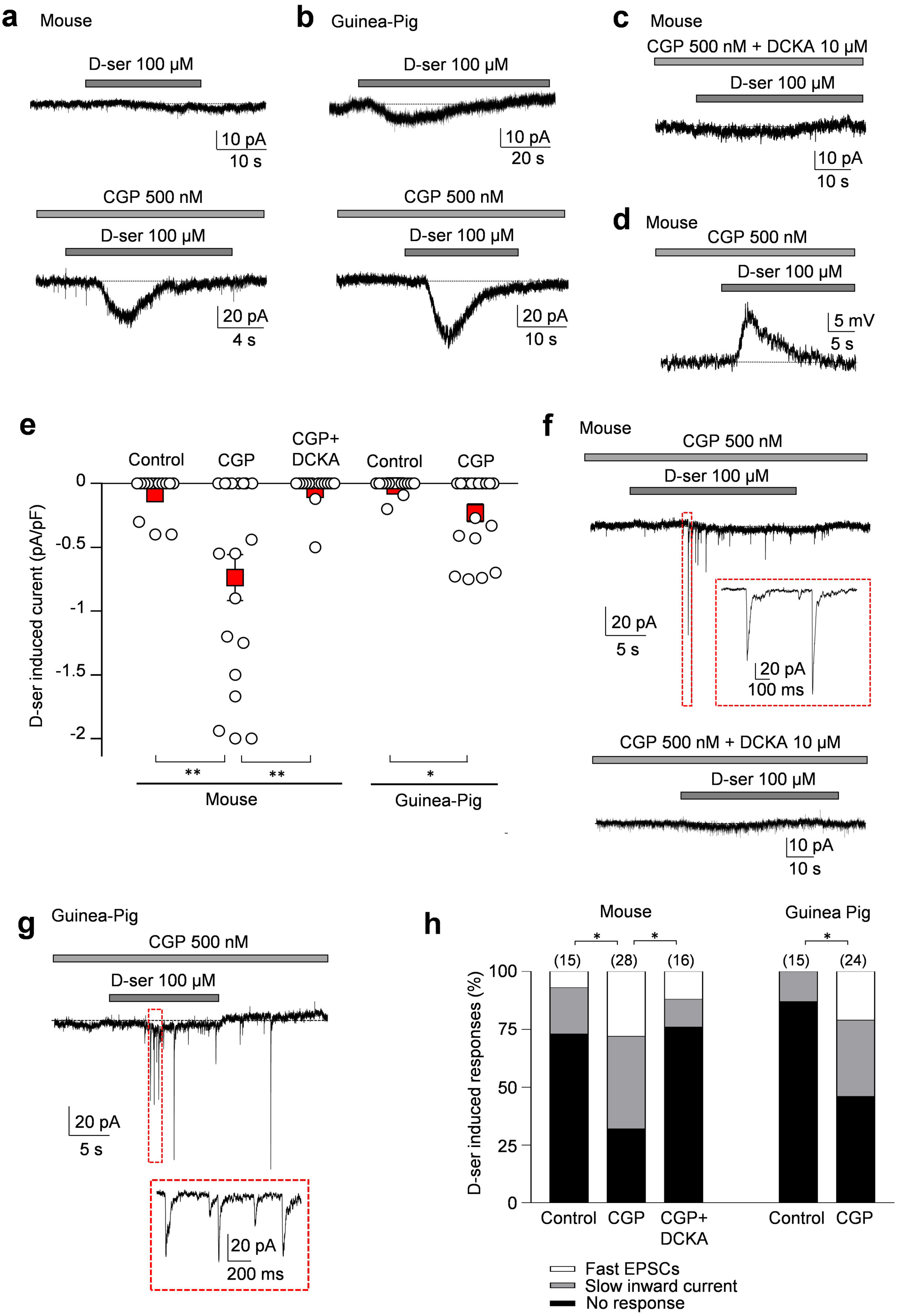
D-Ser induces GluN1-GluN3-like inward current in mouse myenteric neurons. **a, b**) Example traces illustrating D-Ser-evoked currents in mouse (**a**) and guinea-pig (**b**) myenteric neurons recorded *in situ* in control conditions (upper traces) and following preincubation with 500 nM CGP (lower traces). Holding potential -60 mV; CsCl-based pipette solution. **c**) Adding DCKA (10 µM) to the CGP-containing bathing solution abolished D-Ser currents in a mouse duodenal myenteric neuron. Holding potential -60 mV, CsCl-based intrapipette solution. **d**) Example trace illustrating D-Ser-evoked membrane depolarization in a mouse myenteric neuron following preincubation with 500 nM CGP. KCl-based intrapipette solution. **e**) Quantification of D-Ser-induced currents in duodenal mouse and guinea-pig myenteric neurons in control conditions, in the presence of CGP (500 nM) and when DCKA (10 µM) was added to the CGP solution. Mouse: I_Ctrl_ = -0.08 ± 0.04 pA/pF, n = 14; I_CGP_ = -0.74 ± 0.18 pA/pF, n = 19; I_DCKA_ = -0.04 ± 0.04 pA/pF; n = 14. GP: I_ctrl_ = -0.02 ± 0.01 pA/pF, n = 15; I_CGP_ = -0.23 ± 0.07 pA/pF, n = ** P < 0.01; * P < 0.05; Mann-Whitney test. **f**) Example traces illustrating D-Ser-evoked EPSCs in a mouse duodenal myenteric neuron following preincubation with 500 nM CGP or 500 nM CGP + 10 µM DCKA. Holding potential -60 mV, CsCl-based intrapipette solution. **g**) D-Ser-evoked EPSCs in a guinea-pig duodenal myenteric neuron following preincubation with 500 nM CGP. Holding potential -60 mV; CsCl-based intrapipette solution. **h**) Proportion of mouse and guinea-pig myenteric neurons with specified responses to 100 µM D-Ser in control conditions and in the presence of CGP or CGP + DCKA. Number of recorded neurons are indicated. * *P* < 0.05; χ^2^ test.

In few cases, bursts of fast excitatory postsynaptic currents (fEPSCs) occurred in post-synaptic mouse and guinea-pig myenteric neurons during D-Ser application (**Figure 4f,g**). These events, which reflect D-Ser-mediated release of neurotransmitters from presynaptic neurons, were increased in myenteric ganglia treated with CGP78608 both in mice (from 6 to 29%) and guinea pigs (from 0 to 21%). DCKA reversed the effects of CGP78608 on D-Ser-induced fEPSPs in mouse myenteric neurons (**Figure 4f,h**).

### Pharmacological manipulation of enteric GluN1-GluN3 receptors regulates colon contractility

To specify the physiological roles of neuronal GluN1-GluN3 receptor activation on mouse gut motor behavior, we investigated contractility parameters using *in vitro* colonic strips. Longitudinal colonic muscle strips developed spontaneous activity characterized by the occurrence of rhythmic phasic contractions having a mean amplitude of 0.19 ± 0.02 g, a mean frequency of 4.20 ± 0.23 contractions per minute and a mean basal tone of 0.90 ± 0.02 g (**Figure 5**). In line with our patch-clamp recordings, application of D-AP5 (100 µM) had no effects on spontaneous activity (**Figure 5a-c**), confirming the absence of tonically active GluN1-GluN2 NMDARs in colonic contractions under physiological conditions. By contrast, exposure of colonic strips to DCKA (50 μM), which prevents D-Ser binding to both GluN1 and GluN3 subunits and consequently blocks GluN1-GluN3 receptor activity, reduced the area under the curve (AUC) and frequency of ongoing phasic contractions, without affecting the resting basal tone (**Figure 5d-f**). On average, DCKA-induced inhibition of spontaneous colonic contractions reached 15 % when measured over a 5 min post-drug period and could reach 80% at its maximum effect. On the other hand, CGP78608, which acts as a potentiator of GluN1-GluN3 receptor caused a tonic colonic contraction, paralleled by a marked elevation of the basal tonus (**Figure 5g-I, inset in** **Figure 5i**). The phasic contractions were usually ablated during the occurrence of the tonic contraction but resurfaced once the tone began to decrease from the maximum (**Figure 5g,h**). Collectively, these results suggest that ambient D-Ser tunes neuronal GluN1-GluN3 receptor activity and regulates the colonic smooth muscle contractility.

**Figure 5.**
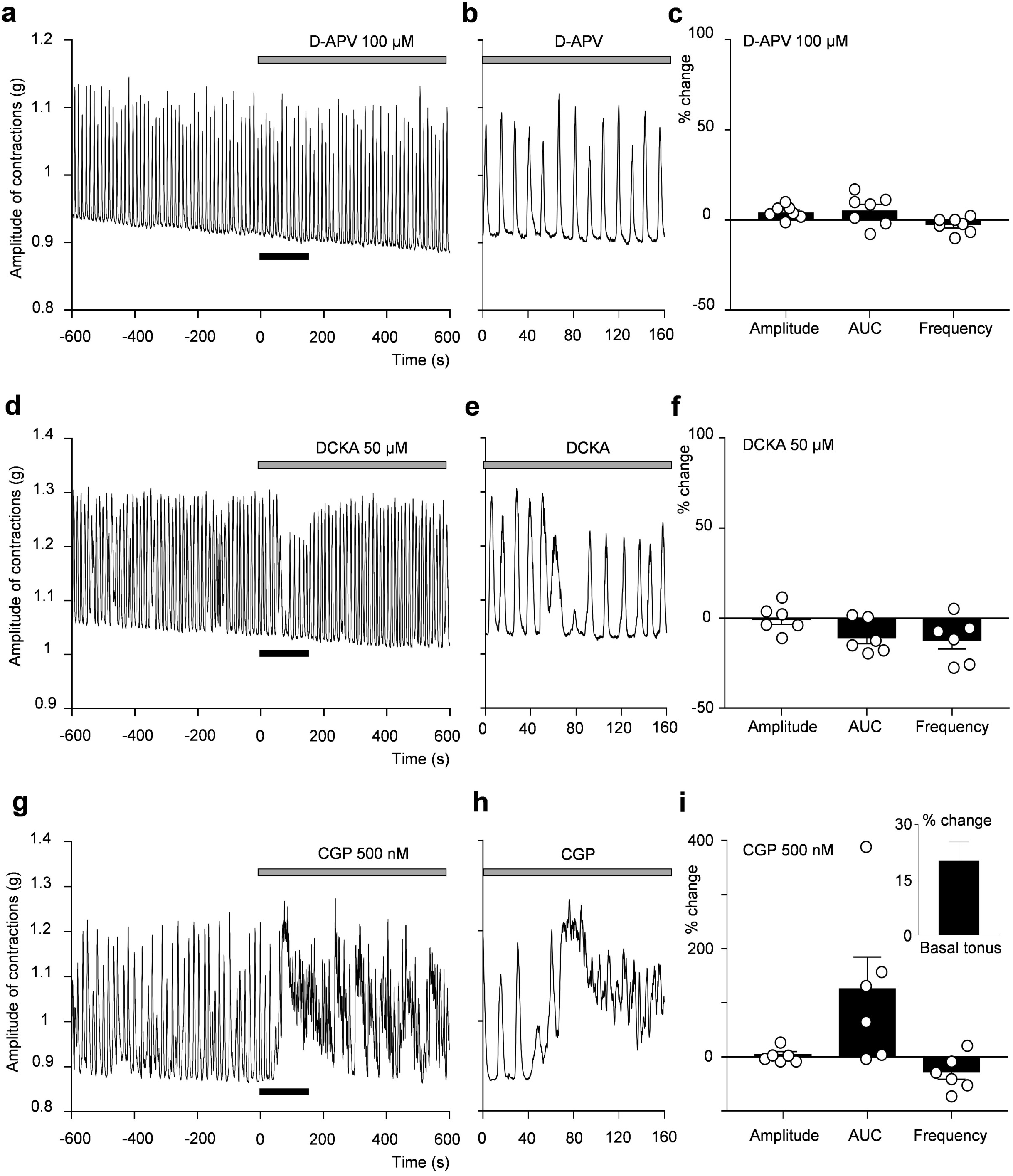
Pharmacological manipulation of GluN1-GluN3 receptors alters motor patterns of isolated strips of mouse colon. DCKA (50 μM, **d-f**) and CGP-78608 (500 nM, **g-i**) have opposite effects on spontaneous contractions of mouse colonic strips. Note that D-AP5 (100 μM, **a-c**) had no effects on colonic contraction, consistent with the lack of GluN2-containing receptors. The rightmost panels illustrate the mean changes in amplitude, area under curve (AUC) and frequency of phasic colonic contractions for individual strips. Data normalization was determined for 5 min-recording periods after (post-drug) and before (pre-drug) drug application, i.e. 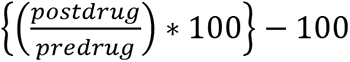. Inset in (**i**) illustrates the mean percentage change of the basal tonus in the 6 strips treated with CGP-78608 (500 nM).

### Genetically-driven loss of D-Ser levels alters gut transit

Given that colonic contractility is impaired upon pharmacologically acting on the GluN1-GluN3 receptor D-Ser binding site, we hypothesized that deletion of SR in mice would impact GI activity. We first performed histopathological analysis on hematoxylin-eosin-stained portion of the whole GI tract of WT and SR^-/-^ mice. Surface area, thickness of the muscularis and submucosa layers, number, and size of the villosities as well as lumen area for the ileum, proximal, medial, and distal parts of the colon were analyzed. We found no obvious clinicopathological differences between WT and SR^-/-^ mice, except for a modest change in the thickness of the muscularis and villi structure in the different segments of SR^-/-^ colon (**Extended data Fig. 5**).

We further compared GI motility of WT and SR^-/-^ mice by measuring fecal pellet output and the time in carmine red expulsion. The stool frequency (119 and 116 pellets/hr for WT and SR^-/-^ mice, respectively) was not significantly different in SR^-/-^ mice compared with WT mice. However, the whole gut transit time was significantly (p = 0.0016) reduced in SR^-/-^ mice to 93.63 ± 5.21 min (n = 8) compared with WT mice (132.14 ± 7.65 min, n = 7) (**Figure 6a**). Consistently, the fecal pellets of SR^-/-^ mice contained significantly more fluid content (**Figure 6b**). Such alterations in GI activity were not resulting from differences in the total length of the GI tract as illustrated in **Figures 6c and 6d**.

**Figure 6.**
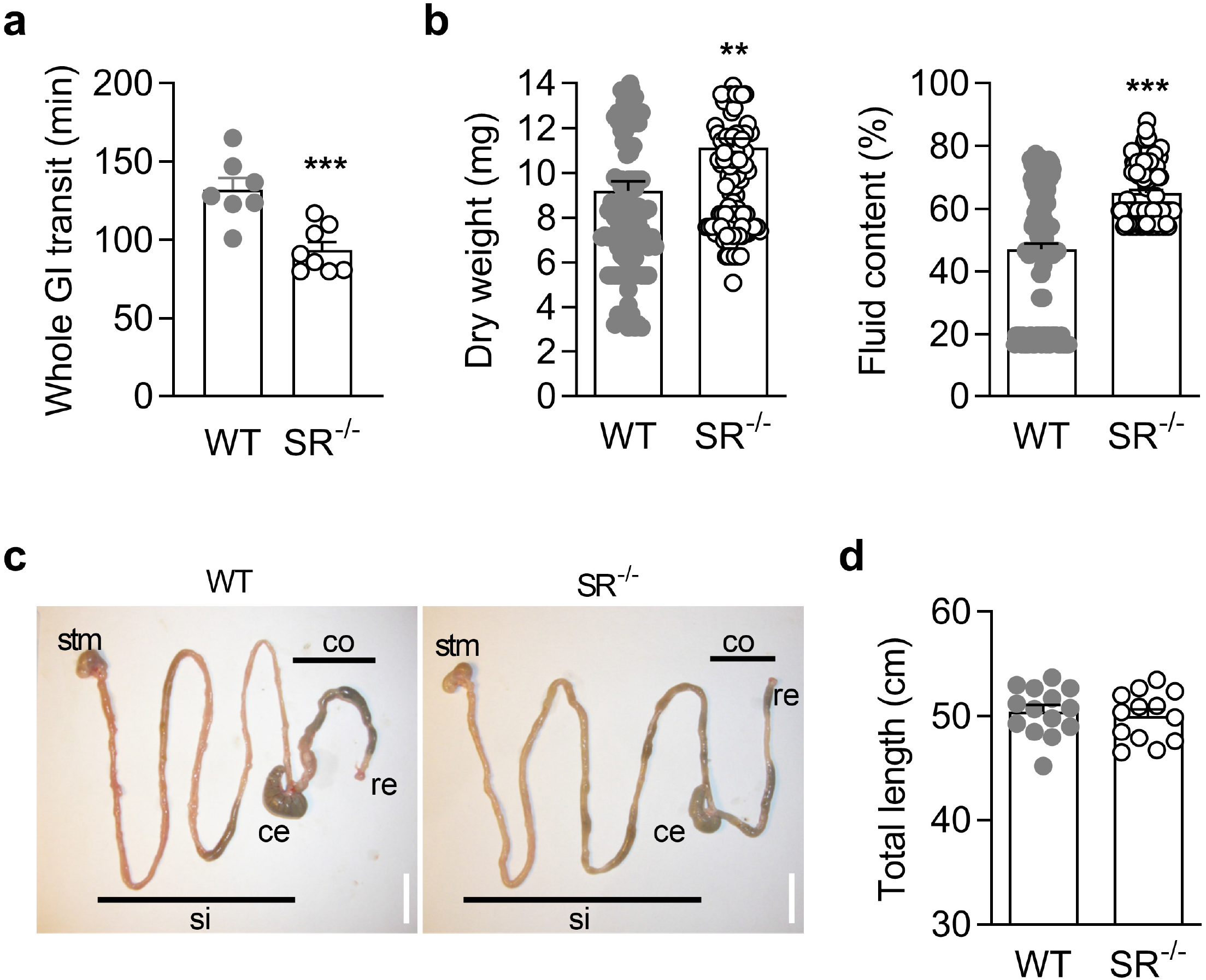
Deletion of SR alters gut transit and pellet output. **a**) Bar graph showing that the duration of the whole GI transit is reduced in SR^-/-^ mice (n = 8) when compared to the littermate wild-type (WT) mice (n = 7). ***p=0.0016 using unpaired t test. **b**) The quality of the pellet output is altered in SR^-/-^ mice. Both the dry weight (left) and fluid content (right) are increased (n = 119 pellets for WT vs 116 for SR^-/-^ mice). **p<0.01, ***p<0.001 using Mann-Whitney test. **c-d**) Comparison of the total GI length between WT (n = 14) and SR^-/-^ (n = 13) mice reveals no obvious changes in the anatomy of the GI from stomach (stm) to the rectum (re). Abbreviations: ce, cecum; co, colon; si, small intestine. P = 0.5584 using Man-Whitney test. **a-d**, errors bars show s.e.m.

## Discussion

Our findings revealed a novel function for D-Ser in the enteric nervous system. We discovered that non-conventional GluN1-GluN3A NMDARs, but not GluN2 containing NMDARs, are physiologically operational in myenteric neurons of mice and guinea-pigs. We provide evidence that ambient D-Ser serves as an agonist at these receptors and thus controls smooth muscle contraction and gut transit. These results add a new active component to the already complex list of neuromodulators and transmitters of the ENS and extend D-serine’s breadth of physiological actions by uncovering its involvement in GI activity.

The exploration of the roles of D-Ser in brain has gained extensive attention during recent years and this atypical neurosignalling molecule is increasingly recognized as a major modulator of synaptic plasticity and cognitive processes through its actions on the NMDARs and GluD glutamate receptors. Both glial and neuronal distributions of D-Ser and SR have been reported in mammalian brains^63, 64^. Although Dai and colleagues discovered that D-Ser produced by SR in the intestine regulates sleep in flies^39^ and several studies have reported the presence of substantial amounts of D-Ser in the rodent gut microbiota^40, 41^, its function in GI tract has remained enigmatic. We report that D-Ser is present at typical concentrations in different segments of the mouse small intestine and the colon as well. D-Ser immunoreactivity was found to be localized in the cell bodies of enteric neurons, but not in glial cells, at least in physiological conditions. Within enteric neurons, D-Ser and SR were commonly encountered in neurons with known markers of IPANs indicating a potential for D-Ser production and secretion from these neurons. Though under-represented, our data also suggests that some nNOS-positive neurons, which are putative inhibitory motoneurons or descending interneurons, also accumulate D-Ser. Metabolic D-Ser degradation is typically achieved by DAAO, a flavin-dependent oxidase widespread to many cells and organs^51^. We found DAAO to be absent in the colon and present only in the proximal part of the mouse small intestine, confirming previous studies^41, 65^, and supporting the absence of any functional effects of CBIO, a known inhibitor^66, 67^, on colonic neurons (data not shown). Intriguingly, we observed a reduction of about 50% in D-Ser amounts in the gut of SR null mutant mice, contrasting with the 80-90% decrease consistently reported in brain D-Ser levels^68, 69^. Our findings thus support that, additional sources, including diet and intestinal gut microbiota bacteria, are likely supplying the gut with D-Ser. Collectively, these data nevertheless indicate that the molecular components for the production and degradation of D-Ser, although sparsely expressed, are present in the ENS and GI tract.

The presence of D-Ser along with GluN1 and GluN2D subunits spurred investigations into a possible functional relationship between them. To be activated, the conventional NMDARs requires simultaneous binding of glutamate to the GluN2 subunit and the co-agonist glycine or D-Ser to GluN1^30–,32^, with the latter being more potent both at binding to the co-agonist site and stimulating the receptor. Despite extensive investigation in mouse and guinea-pig myenteric neurons, we failed to detect functional GluN1-GluN2 NMDARs both at post- and pre-synaptic levels, ruling out a contribution of these receptors in D-Ser action. Consistently, D-AP5 failed to alter the spontaneous activity of mouse colonic strips. This observation validates the lack of any physiological contribution of GluN1-GluN2 NMDARs in the activity of the colon, which however does not preclude a possible role for these receptors in pathological conditions. Previous studies have produced conflicting data regarding the presence of these conventional NMDARs in enteric neurons^13, 17, 26^. While Liu et al.^13^ reported that D-AP5 inhibits glutamate-induced depolarization in enteric neurons, others observed no effects of the NMDAR antagonists, MK-801 or D-AP5, on glutamate-induced excitation or calcium transients^17, 26^. Our *in situ* patch clamp recordings indicate that mouse and guinea-pig myenteric neurons do not respond to NMDA exposure under physiological conditions, yet they exhibit fast developing inward currents in response to L-glutamate. Therefore, glutamatergic responses are likely to relate to the presence of other glutamate receptors such as Kainate or AMPA receptors^70^.

Our findings point towards the contribution of GluN3-containing receptors. GluN3A and GluN3B can associate with multiple NMDAR subunits, however, only tri-heteromeric GluN1-GluN2-GluN3 and di-heteromeric GluN1-GluN3 assemblies produce surface expressed receptors in heterologous cell systems. The tri-heteromeric GluN1-GluN2-GluN3 complexes respond to NMDA and glutamate and are thus not at play in enteric neurons. Diheteromeric GluN1-GluN3 complexes are not activated by NMDA or glutamate, are insensitive to D-AP5 and Mg^2+^ and function as excitatory glycine-gated receptors^71, 72^.

Our data are consistent with the involvement of GluN1-GluN3 receptors, i.e. i) activation by D-Ser alone in the absence of glutamate or NMDA, ii) insensitivity to the GluN2 glutamate binding-site antagonist D-AP5^62, 71, 72^ and iii) inhibition by the dual GluN1 and GluN3 glycine-binding site antagonist DCKA. Furthermore, the GluN1 antagonist CGP78608, which unmasks GluN1-GluN3A receptors^62, 71^ potentiated D-Ser signals both in patch clamp recordings and calcium imaging. Our findings align with single cell RNA-sequencing assays, which identified expression of GluN1 (*Grin1*) and GluN3 (*Grin3a*) subunits, along with unique electrophysiological channels (Kcnn3, Ano2, Scn11a) in mouse IPANs^59, 73^. Altogether, these results provide the first demonstration that D-Ser excitatory GluN1-GluN3 receptors are functional in enteric neurons in contrast to data in the CNS where glycine is the main co-agonist of these receptors^71^.

A teasing remaining question then is whether ambient D-Ser modulates GluN1-GluN3 NMDAR functions *in vivo*. Our data show that D-Ser-induced currents in *ex-vivo* LMMPs are barely detectable in control conditions but consistently turn into more prominent responses in the presence of CGP78608. This observation suggests that the main fraction of GluN1-GluN3 NMDARs exists in a D-Ser-bound inhibited/desensitized state. Accordingly, acute application of D-Ser only elicited calcium transients in a fraction of neurons, but this effect was potentiated by CGP78608, as would be expected from GluN1’s inhibition relief of D-Ser-bound GluN1-GluN3 receptors. Because D-Ser binds to the GluN3A subunit with a ∼10-fold higher affinity than GluN1, we suggest that D-Ser concentration in LMMPs is high enough to maintain those receptors silent, consistent with the high level of D-Ser found in LMMPs and gut tissues. Hence, only a subset of GluN1-GluN3 NMDARs is available for activation in LMMPs. Yet, the final concentration of D-Ser that reaches the target effector cells is still unknown *in vivo*. Reduction of D-Ser levels by ∼50% in SR^-/-^ mice was found to speed-up overall GI-transit time *in vivo*, indicating that D-Ser through GluN3 receptors may be important in controlling small intestine motility. In addition, SR^-/-^ mice have loose stools, indicating impaired absorptive function, which is expected from a change in GI-transit that deprives the fecal material of adequate exposure to mucosa. Collectively these phenotypic alterations are in line with the idea that GluN1-GluN3 receptors in the ENS coordinate muscle movements underlying propulsion of content and absorption. However, it is also possible that changes in the activity of the extrinsic innervation of the GI tract, arising from both the sympathetic and parasympathetic branches of the autonomic nervous system, may contribute to the observed effects in vivo. At the organ level, investigation of contractility parameters using *in vitro* colonic strips suggests that activation of GluN1-GluN3 NMDARs acts on muscle contraction, as DCKA reduces contraction amplitude while CGP78608 has an opposed effect. Thus, GluN1-GluN3 NMDARs in the colon are important for normal colorectal motility. Because the absorption rate in the colon is slow compared to other GI segments (due to tight mucosa), an important requirement of colonic motility is to propel the luminal content slowly to prolong exposure to the mucosa. Thus, variation in the amount of D-Ser may lead to colonic dysmotility leading to constipation or diarrhea.

Decoding the role of GluN1-GluN3 NMDAR and D-Ser in ENS remains exceptionally challenging, owing to its complex organization, particularly with regards to its heterogeneous population of enteric neurons that innervate different targets and interactions with the microbiome and immune system. Collectively, our data suggest that D-Ser produced by enteric neurons may act at GluN1-GluN3 NMDARs to regulate gut neural pathways underlying peristalsis and the balance between transit and absorption. Since the ENS and gut microbiota reside in close anatomical proximity, microbiota-derived D-Ser might also modulate GluN1-GluN3 NMDARs and GI functions. Indeed, gut motility is highly dependent on microbiota^74^ and different studies have demonstrated that several commensal bacteria produce D-Ser^40^. Because the gut epithelial barrier expresses transporters for AAs including D-Ser, it is tempting to imagine that part of the microbiota-derived D-Ser can reach the enteric GluN1-GluN3 receptors. Such crosstalk between microbiota and the ENS metabolism implying D-Ser is offering an appealing framework for tailoring new strategies for treat or prevent GI diseases by targeting D-Ser pathway.

## Methods

### Animals

All experiments were conducted in accordance with European and French directives on animal experimentation and were approved by the CNRS, Aix-Marseille University and KU Leuven Animal Ethics committees (2010/63/UE). Animals were bred in-house in polycarbonate cages and maintained on a 12:12-h light/dark cycle in a temperature (22°C) and humidity-controlled room. Animals were given access to food and water *ad libitum* except during food fasting period preceding some experiments. Age-matched (12–16-week-old) SR knock-out (SR^-/-^) mice and wild-type (WT) mice on a C57BL/6 background were used. Tricolor strain guinea pigs weighing 250-400 g from the inbred colony (Marseille, France) were also used. Mice and guinea pigs were killed before experiments by cervical dislocation or decapitation after isoflurane anesthesia.

### Determination of D- and L-AAs levels by capillary electrophoresis with laser induced fluorescence (CE-LIF)

Luminal contents and muscularis layers from the small intestine from WT and SR^-/-^ mice were isolated following the protocols described earlier^41^ with slight modifications. Middle segments (2-3 cm) of the ileum and colon were freshly dissected. The inner part of each piece was washed out 3 times with 2 mL of ice-cold sterile PBS, resulting in 6 mL of washout referred to as ‘luminal content’ or ‘lumen’. Then tissue pieces were opened longitudinally, and the mucosa and the submucosa were dissected away. The remaining tissue pieces referred to as ‘muscularis layers’ or ‘wall’ were dried and placed in cryogenic tubes filled with ice-cold methanol. Samples were kept at -80°C until used for analysis. Each luminal content tube was supplemented with 500 μL methanol and then dried using a SpeedVac system. The muscularis layers were homogenized, and the supernatants were collected and dried using a SpeedVac. The pellets were sequentially rinsed with 1 mL acetonitrile, 1 ml 80:20 methanol: H_2_O, and 500 μL of water. Each of these solutions was combined with the original dried supernatant and evaporated to dryness using a Speedvac to finally give the tissue extract samples. All samples were reconstituted in water and then derivatized before determination of D- and L-AAs levels using capillary electrophoresis with CE-laser induced fluorescence (LIF).

Naphthalene-2,3-dicarboxaldehyde (NDA) derivatization for LIF detection was accomplished by mixing samples and D-Ser standards with potassium cyanide (20 mM KCN in 100 mM borate buffer) and NDA (20 mM in acetonitrile) in a 1:4:4 ratio. Following derivatization, the reaction mixture was diluted in water and was analyzed by CE-LIF. To confirm the presence of D-Ser, gut samples were treated with D-amino acid oxidase from porcine kidney (*Pk*DAAO) (Catalog# A5222, Sigma-Aldrich, St. louis, MO). The sample was mixed with *Pk*DAAO (15 U/mL), flavin adenine dinucleotide (5 mM), catalase (68 μg/mL) from bovine liver, and PBS 1X in a 1:2:1:1:5 volume ratio. The mixture was incubated in a water bath at 37 °C for 6 h. The enzyme-treated sample was dried and reconstituted in 1 μL water prior to derivatization and analysis by CE-LIF.

Separations were performed using a PA 800 plus Pharmaceutical Analysis System equipped with LIF detection (AB SCIEX, Framingham, MA). The laser emitting wavelength of 440 ± 8 nm was operated at 3 mW. A band-pass filter of 490 ± 15 nm was used to only permit the fluorescence emission band for detection. Bare fused-silica capillary was used for all separations. The capillary had the total/effective lengths of 40/30 cm with inner/outer diameters of 50/360 μm. The separation buffer with final concentrations of 62 mM 2-(N-morpholino)ethanesulfonic acid (MES) buffer (pH 6), 7 mM potassium bromide, and 330 ppm quaternary ammonium β-cyclodextrin solution was prepared daily into water. For D-Ser separation, 10 kV of reverse polarity at 20 °C was applied. The standard calibration curve was generated to quantitate D-Ser in the gut samples based on the analyte peak area. Protein quantification was performed using the Micro BCA Protein Assay Kit (ThermoFisher, Walham, MA)to further compare amino acids levels between samples.

### Reverse Transcription Polymerase Chain Reaction (RT-PCR)

Total RNA was extracted from brain and colon LMMP preparations of WT and SR^-/-^ mice with TRIzol reagent (Ambion) according to the manufacturer’s guidelines. Reverse transcription reactions with oligo(dT)20 primers were performed with the Superscript III reverse transcriptase (Invitrogen) following the manufacturer’s protocol (Invitrogen) using 3 µg of total RNA. PCR on reverse-transcribed cDNA was performed with specific primers to mouse transcripts. Sequences of primers and expected sizes of the amplicons were described in the **Supplementary Table 1**. PCR products were separated on a 2% agarose gel.

### Western Blot analysis

Brains and colonic LMMP preparations isolated from mice or guinea pigs were homogenized by sonication in ice-cold lysis buffer (Tris 50 mM, pH 6.8 containing SDS 0.5%, EDTA 2 mM, PMSF + Complete EDTA free protease inhibitors cocktail (Roche Diagnostics) w:v = 1:10) and centrifuged 10 min at 1,500g to remove the crude nuclear pellet. Protein extracts (20 µg for brain and 40 μg for LMMP per lane) were subjected to electrophoresis (10% (vol/vol) SDS-polyacrylamide gel) and electroblotted onto nitrocellulose membranes (0.2 μm; Pall Life Science). Immunodetection was accomplished by enhanced chemiluminescence (ECL) using the ECL Prime Western Blotting Detection Reagent (Amersham Biosciences). Molecular sizes were estimated by separating prestained molecular weight markers (10.5–175 kDa) in parallel (Nippon Genetics). Primary and secondary antibodies and their dilutions are listed in **Supplementary Table 2**. Digital images of the chemiluminescent blots were captured using the GBOX system (Syngene) with the GeneSys image acquisition software.

For DAAO immunoblots, two 6-week-old mice were anesthetized and perfused with ice-cold PBS before dissection of intestinal tissue. The protocol for isolation of intestinal epithelial layer was described previously^41^. Non-epithelial tissue was rinsed in PBS twice. Tissue was stored in –80°C until use. Epithelial and non-epithelial tissue was homogenized in a lysis buffer (150 mM sodium chloride, 1.0% NP-40, 50 mM Tris (pH 8.0), and a protease inhibitor cocktail, Complete EDTA-free (Roche, Basel, Switzerland)) and spun down at 15,000g at 4°C for 5 min. Supernatants were subjected to SDS-PAGE and protein was transferred to nitrocellulose membranes. Blots were blocked in 10% skim milk in PBS with 0.1% Tween-20 (PBST) with constant shaking for 1 hour. The membranes were rinsed in PBST and immersed in PBST with a rabbit polyclonal antibody to mouse DAAO^41^ or a rabbit monoclonal antibody to GAPDH at 4°C overnight. Membranes were subsequently rinsed in PBST, then incubated with an appropriate secondary antibody conjugated with horseradish peroxidase for 1 hour. The membranes were rinsed in PBST and bound antibodies detected with ECL (GE Healthcare Life Sciences, Chicago, IL, USA).

LMMP preparations isolated from human colons were placed into RIPA buffer (0.5 M Tris-HCl, pH 7.4, 1.5 M NaCl, 2.5% deoxycholic acid, 10% NP-40, 10 mM EDTA), containing a cocktail of protease inhibitors (Roche Diagnostics), snap frozen on dry ice and homogenized. Protein concentration was measured using a BCA protein assay kit (Bio-Rad, Hercules, CA). Aliquots containing 40 μg of proteins were resolved in 8% SDS-PAGE under reducing conditions and transferred to a nitrocellulose membrane. The membrane was blocked with 1% BSA in PBS + 0.05% Tween-20 for 1 hour at room temperature and incubated with primary antibodies overnight at 4°C. After washing, the membrane was incubated with appropriate horseradish peroxidase (HRP) conjugated secondary antibody diluted in PBST milk before imaging.

### Immunohistochemistry and confocal microscopy analysis

The LMMP preparation has been detailed previously^57, 59^. In brief, a 2 cm segment of each intestine portion was removed from animal, placed in a Sylgard coated dish containing oxygenated standard Krebs solution (in mM: 118 NaCl, 4.8 KCl, 1 NaH2PO_4_, 25 NaHCO_3_, 1.2 MgCl_2_, 2.5 CaCl_2_, and 11 glucose, equilibrated with 95% O_2_-5% CO_2_, pH 7.4 and supplemented with 2 µM atropine and 6 µM nicardipine). The myenteric plexus was exposed by dissecting away the mucosa, the submucosal plexus, and the circular muscle layers. LMMPs were then fixed by immersion for 1-2hr in PBS containing 4% paraformaldehyde (complemented with 0.5-1% glutaraldehyde for D-serine immunolabelling). Whole-mount preparations were subjected to immunohistochemistry as described previously^75^. Antibody details are supplied in **Supplementary Table 2**. Images were acquired through the 40X objective (PlanFluor, 0.75 NA) of an upright Leica confocal laser scanning microscope. No staining was observed in SR^-/-^ mice or when the anti-D-Ser antibody was incubated in the presence of liquid conjugate against D-Ser. The confocal images were acquired using the Leica TCS software with a sequential mode to avoid interference between each channel and without saturation of any pixel. Moreover, emission windows were fixed for each fluorophore in conditions where no signal is detected from the other fluorophore. Stack images were taken, and a Z-projection was made with ImageJ software (http://rsb.info.nih.gov/ij) in average projection mode.

### Histochemistry for DAAO

Activity-based DAAO labeling was performed using a previously reported method^41^. After deep anesthesia, 6-week-old mice were perfused transcardially with 2% paraformaldehyde in PBS. Tissues were dissected, cryoprotected and 15 µm sections were made on a cryostat (Leica, Wetzlar, Germany). Sections were rinsed in PBS and immersed in DAAO-activity staining solution as defined previously^41^. Fluorescence signals were visualized using a BZ-9000 fluorescent microscope (Keyence, Osaka, Japan). Each section being compared was imaged under identical conditions.

### Histopathological analysis of SR^-/-^ mouse gut

We favor the so-called gut bundling technique as it provides high quality cross sections through villi and crypts and enable quantification of cell features or villus/crypt lengths. The entire GI was extracted from fasted mice (6 WT and 6 SR^-/-^). The lumen was gently flushed with ice-cold PBS to remove any fecal and digesta content. The large and small intestine was cut to isolate the medial part of the ileum (48.5-51.5%), the proximal (10-12.5%), medial (48.5-51.5%) and distal (77.5-80%) parts of the colon. Gut segments were then placed overnight with 10X excess of tissue volume 4% paraformaldehyde fixative and thereafter transferred to tissue cassette to be embedded in paraffin. Cross sections (10 μm) were obtained using a cryostat prior to be floated onto glass slides. Sections from WT and SR^-/-^ mice were simultaneously colored with hematoxylin/eosin/safran (HES; Merck, Darmstadt, Germany). Once stained, slides were scanned and proceeded for morphometric analyses. Villus size (height and width), lumen and total sections areas, thickness of both smooth muscle layer and submucosa layer were scored from at least 10 sections per mice. Villus height was measured from the basal layer of the submucosa to the ending of the villus in the ileum and colon.

### Calcium Imaging

Colon segments (approximately 5 cm long) were collected and immersed in 95% O_2_/5% CO_2_ -bubbled Krebs solution (in mM: 120.9 NaCl; 5.9 KCl: 1.2 MgCl_2_; 1.2 m NaH_2_PO_4_; 14.4 NaHCO_3_; 11.5 Glucose; 2.5 CaCl_2_). Tissues were opened along the mesenteric border and pinned flat with mucosa up in a silicone elastomer-coated dish. After removing the mucosa and submucosa, the tissues were flipped over and the longitudinal muscle layer was removed to expose the myenteric plexus. These preparations (up to 5 per colonic segment) were stabilized over an inox ring using a matched o-ring^76^. The ring preparations were loaded for 20 min in bubbled Krebs containing 0.01% (vol/vol) Cremaphor and 1 μM Fluo-4-AM. After a 20 min washout period the tissues were transferred to a coverglass chamber mounted on the microscope stage. 3D recordings of the Fluo-4 signals were made on an inverted spinning disk confocal microscope (Nikon Ti-Andor Revolution – Yokogawa CSU-X1 Spinning Disk (Andor, Belfast, UK) with a Nikon 20x lens (NA 0.8), excitation 488 nm and detection 525/50 nm. A Piezo Z Stage controller (PI) was used to record fast 3D stacks at 1Hz. For other sets of experiments, 2D recordings were made on an inverted Axiovert 200M microscope (Carl Zeiss, Oberkochen, Germany), 20x 0.8 NA objective lens, equipped with a monochromator (Poly V) and cooled CCD camera (Imago QE), both from TILL Photonics (Gräfelfing, Germany). Fluo-4 was excited at 470 nm (50 ms exposure) and images were acquired at 520/50 nm at a 2 Hz framerate. The tissues were constantly supplied with 95% O_2_ / 5% CO_2_-bubbled Krebs at room temperature via a gravity fed perfusion system (1 ml/min). The pipette tip, in which 5 perfusion lines converged, was positioned just above (250 µm) the ganglion of interest. A magnetic valve system allowed instantaneous switching between control and drug-containing Krebs solutions. For CGP78608, a preincubation (15 min) was used before preparations were transferred to the recording microscope. A 75 mM K^+^ depolarization (5 s) was used to depolarize the ganglion and identify all Fluo4 labelled cells. All solutions contained 2 µM nifedipine (N7634, Sigma-Aldrich) to minimize spontaneous movement.

All image analysis was performed with custom-written routines^77^ in Igor Pro 8 (Wavemetrics, Lake Oswego, OR, USA). 3D image sequences were imported, and maximum projections were generated from the image planes that contained the myenteric plexus. These 2D images were analysed in the same way as the images recorded on the 2D widefield system. First, regions of interest were drawn in the images to extract the temporal information from individual cells. Peak detection and peak amplitude calculation were performed using custom written procedures as previously described^82^. The average Ca^2+^-signal intensity was calculated, normalized to the initial Fluo-4 values, and reported as F/F0. Responses were considered when the [Ca^2+^]_i_ signal increased above baseline by at least five times the intrinsic noise. [Ca^2+^]_i_ peaks were calculated for each response, with the peak amplitude taken as the maximum increase in [Ca^2+^]_i_ from baseline (Fi /F0). The total number of cells responding to each drug exposure was counted and expressed as a percentage of the control.

### Whole-cell patch-clamp recordings from intact myenteric ganglia

Whole-cell patch-clamp recordings were conducted at room temperature on LMMP preparations obtained from duodenum and colon of both guinea pigs and mice. The LMMP preparation of gut segment has been detailed previously^57^. In brief, a 2 cm segment of duodenum or colon was removed. The tissue was opened along the mesentery line and pinned mucosal side up on the silicon elastomer basis (Sylgard; Dow Corning) of a recording dish containing oxygenated standard Krebs’ solution. The myenteric plexus was exposed by dissecting away the mucosa, the submucosal plexus, and the circular muscle layer. The recording dish was then placed on an inverted microscope stage and the LMMP preparation was continuously superfused with Krebs’ solution gassed with 95% O_2_-5% CO_2_. The upper surface of a ganglion was exposed to 0.02% protease type XIV (Sigma) for 3–5 min and the surface of neurons was cleaned by sweeping over the ganglion with a hair fixed at the tip of a microelectrode. The standard Krebs’ solution used to bath the LMMP preparation consisted of the following (in mM): 118 NaCl, 4.8 KCl, 1 NaH_2_PO_4_, 25 NaHCO_3_, 1.2 MgCl_2_, 2.5 CaCl_2_ and 11 glucose and was equilibrated with 95% O_2_-5% CO_2_, pH 7.4. Atropine (2 µM) and nicardipine (6 µM) were present throughout the experiments to prevent spontaneous muscle movement. Drugs, diluted into the standard Krebs’ solution (see above), were locally applied via a gravity perfusion system directly onto the myenteric ganglion under study. Patch pipettes were pulled from thick-walled borosilicate glass capillaries (Harvard Apparatus) and had a resistance of 3-5 MΩ. The intracellular solution used to record in the voltage-clamp mode contained (in mM): 140 CsCl, 4 NaCl, 1 CaCl_2_, 2 MgCl_2_, 2 EGTA, 10 HEPES. For current-clamp recordings, the intrapipette solution consisted of (in mM): 140 KCl, 4 NaCl, 1 CaCl_2_, 2 MgCl_2_, 2 EGTA, 10 HEPES. pH was adjusted to 7.35 with CsOH or KOH (∼300 mOsm/L).

To elicit EPSPs in enteric neurons, a concentric bipolar electrode (FHC, Bowdoin, USA) was placed over one of the interganglionic bundles lying circumferentially to the recorded ganglion. Nerves fibers were electrically stimulated using 1 ms, 0.5- to 2-mA constant current pulses delivered every 15 s from an ISO-flex stimulus isolation unit driven by a Master-8 pulse stimulator (A.M.P.I., Israel).

All recordings were made using an Axopatch 200B amplifier (Molecular Devices), low-pass filtered at 2 kHz, and digitized at 25 kHz. The series resistance measured in the whole-cell mode was 5-8 MΩ and was compensated by 70-80%.

### Spontaneous contractions of isolated proximal colon from mice

The entire colon was removed and placed into Krebs’ buffer at room temperature. The colon was cleaned from connective tissues and the proximal part (2 cm from the ileocolonic junction) was cut along in the longitudinal direction. Strips (8-10 mm long; 2-3 mm large) were tied with silk threads to secure them between the holder and the isometric transducer (it-1, EMKA Technologies, France) and mounted under 10 mN initial tension in 5 mL organ baths (EMKA Technologies, France) containing Krebs-Henseleit solution (composition in mM: NaCl 114, KCl 4.7, CaCl_2_ 2.5, MgSO_4_ 1.2, KH_2_PO_4_ 1.2, NaHCO_3_ 25, glucose 11.7; equilibrated with 95% O_2_-5% CO_2_, pH 7.4) and kept at 37°C.

Contractile responses were measured using isometric tension transducers connected to amplifiers and to a data acquisition system (PowerLab 8/35, ADInstruments Pty Ltd., Castle Hill, Australia). Colonic tissues were allowed to equilibrate for 60 minutes during which the Krebs’ solution was replaced every 10-15 min. In strips demonstrating spontaneous regular contractions, after a 30-min recording period, D-AP5 (100 µM), CGP78608 (500 nM) or DCKA (50 μM) were added for 15 to 30-min periods. Then, tissues were washed at least three times during an additional 15-min period before the addition of KCl (80 mM) to test tissue viability. The basal tone was determined as mean tension between the peaks of contraction before drugs addition.

### Whole-gut transit time and pellet production

Whole GI transit duration was measured with carmine red simultaneously in WT and SR^-/-^ mice after fasting and according to the procedure described^78, 79^. Carmine red was prepared as 1% solution with 5% sucrose (Sigma-Aldrich) in 2% agar. Mice were individually housed and fasted before the test. In the morning, 6 carmine red agarose pieces were given to mice. T0 refers to the time when mice started to eat. When mice had eaten the totality of carmine red pieces, they were placed on iron gate cage and monitored every 15 min to see the appearance of the first carmine red feces. The whole-gut transit time was defined as the interval between T0 and the time of first observance of carmine red feces. Fecal pellets excreted from WT (n=119) and SR^-/-^ (n=136) mice were collected at zeitgeber +4 and weighed (ie wet weight). Feces were next dried 24h at 37°C and weighed again, which represents weight of dry feces and fluid content was then calculated as described^85^.

### Drugs, enzymes and solutions

Appropriate stock solutions were diluted to the final concentration with perfusing media containing vehicle (<1/1000) just before application. Controls were made with vehicle to ascertain any side effects. Salts were all from Sigma (St Louis, MO, USA). Tricine, L-glutamate, Kainate, Salicylate and D-Ser were purchased from Sigma. N-Methyl D-aspartate (NMDA), CGP78608, 5,7-Dichlorokynurenic acid (5,7DCKA), CNQX and D(−)-2-Amino-5-phosphonopentanoic acid (D-AP5) were from Tocris Biosciences. Tetrodotoxin (TTX) was from Alomone Laboratories (Jerusalem, Israël). All drugs were stored as aliquots at −20°C and were diluted in Krebs to achieve the desired final concentration.

### Data Analysis

Quantitative data are expressed as mean ± standard error of the mean (s.e.m.), or otherwise stated. Graphics and statistical analyses were performed using GraphPad Prism (GraphPad Software, San Diego, CA, USA) excepted for CE-LIF for which Origin 2018 software was used. Data were tested for normality using the Shapiro-Wilks test or D’Agostino and Pearson omnibus test. Outliers were identified by Grubb’s test. Normally distributed data were compared by Student’s t-test for paired or unpaired values. Data not normally distributed were compared by Mann-Whitney test or Wilcoxon’s test. Difference was considered significant if p < 0.05.

## Acknowledgements

The authors thank Bertrand Coste for his critical reading and feedback on the manuscript. The authors would like to thank Solomon H. Snyder and Joseph T. Coyle for providing SR-null mutant mice. We thank Nadine Clerc for collecting some gut tissues, Nicolas Bandone and Philippe Dales for their help in cryosectionning and HES coloration. This work was supported by the Association Française contre les Myopathies (MJN2 2006-12203 to JPM), Fondation pour la Recherche Médicale (ING20121226166 to JPM; DEQ20130326482 to PD), by recurrent operating grants from CNRS, INSERM, Aix-Marseille University and University of Bordeaux (to PD and JPM). P. Vanden Berghe received support from FWO G0921.15, G0938.18, KU Leuven BOF Methusalem/14/05 and Hercules Foundation AKUL/09/50 to purchase the Andor Revolution Spinning Disk System. JVS received support through the USA National Institutes of Health through P30DA018310 and R01NS031609. Magalie Martineau was a recipient of a PhD fellowship from the National Ministry of Education, Research, and Technology.

## Author contributions

Nancy Osorio, Magalie Martineau: conducted experiments, analyzed and interpreted data, and wrote the manuscript. Amit V. Patel, Werend Boesmans, Vivien Labat-Gest, Virginie Penalba, Grégoire Mondielli, Sandrine Conrod, Amandine Papin, Malvyne Rolli-Derkinderen, Cindy J. Lee, Céline Rouget, Marina Fortea: conducted experiments. Jumpei Sasabe, Jonathan V. Sweedler, Pieter Vanden Berghe: designed experiments, analyzed and interpreted data. Patrick Delmas, Jean-Pierre Mothet: designed and supervised the study, analyzed, and interpreted data and wrote the paper. All authors have critically revised the manuscript and approve the final version before submission.

## Extended data

**Extended data Figure 1. Cell-type distribution of serine racemase and D-Ser in the myenteric plexus.** Gray scale images of splitted channels corresponding to double immunostainings illustrated in Fig. 1. Immunostaining for D-Ser (left) and HuC/D (right), a neuronal pan marker. Scale: 50 µm. **b**) Immunostaining for D-Ser (left) and S100B (right), a glial marker. Scale: 50 µm, **c**) Immunostaining for D-Ser (left) and NeuN (right), a marker of gut sensory neurons. Scale: 50 µm. **d**) Immunostaining for D-ser (left) and neuronal nitric oxide synthase, nNOS (right), a marker of inhibitory motoneurons. Scale: 50 µm. **e**) Immunostaining for SR (left) and NeuN (right). Scale: 50 µm.

**Extended data Figure 2. The D-Ser degrading enzyme DAAO is not present in the distal small intestine and the colon. a**) First lane: Cross-sections showing activity based DAAO labeling. The labeling reveals the presence of DAAO in the proximal s.i, but not in the distal s.i and the colon of WT mouse. Second lane: Cross-sections revealing DAAO activity and counterstained for phalloïdin and DAPI. DAAO is only present in the mucosa layer. Scale: 200 µm. **b**) Zooms into ROI shown in (**a**) for each segment of the GI tractus. Scale: 200 µm. **c**) Western blot showing differential expression of DAAO in mouse small intestine and the colon. Tissues extracts were subjected to electrophoresis and to immunoblotting DAAO and GAPDH. A cerebellar lysate was used as a positive control (pc).

**Extended data Figure 3. The D-Ser degrading enzyme DAAO is not detected in the colon by RT-PCR.** Lack of DAAO transcripts in mouse colon as shown by RT-PCR. (-) indicates no RT. Brain tissue is used as positive control.

**Extended data Figure 4. Molecular profiling of glutamatergic receptors expressed in the myenteric plexus of the colon. a**) RT-PCR performed from LMMPs of mouse colon for the different transcripts. Note that transcripts for the major glutamatergic receptor subunits are present (NMDA receptors: GluN1, GluN2D, GluN3A; AMPA receptors: GluA2 and GluD1-2). When transcripts were not positively detected (GluN2A, GluN2B, GluN2C and GluN3B) in the colonic LMMP brain extracts were shown as positive controls (not shown when amplicons from mouse colon LMMP were detected). cDNAs isolated from mouse colon LMMP and brain were amplified using primers GluN1, GluN2A-D, GluN3A-B, Grid1-2 and GluR2 (see **Extended data Table 1**). Contamination from genomic DNA was routinely tested by omitting the reverse transcriptase in the templates (-). M, 100 bp-ladder DNA size standard. **b**) Immunoblots analysis on colonic LMMPs and brain extracts collected from WT and SR^-/-^ mice. GluN1 N ter but not GluN1 C-ter is present in colon confirming previous report^85^. Immunoblots show positive signals for GluN2C/D, GluN3A-B, GluA2 in the colon of WT and SR^-/-^ (-/-) mice and faint or no detection for GluN2B, GluN2A, GluD1-2 and the vesicular transport type 1 (vGlut1), which is the primary vesicular transporter for glutamate in the CNS. All sample loadings were normalized to actin.

**Extended data Figure 5. Histopathological analysis of GI wall in wild-type and SR^-/-^ mice. a**) Schematic representation of the GI tractus highlighting the ileum and colon segments collected for the analysis. **b)** Cross-sections from WT and SR^-/-^ mice and colored for hematoxylin/phloxine safran are illustrated for each segment. **c**) Schematic representation of a cross-section and the different parameters used in the analysis. **d-i**) Bar graphs summarizing the analysis for each segment of the GI tractus in WT and SR^-/-^ mice. In panels **d** & **e**, N=7-8 sections. In panels **f** & **g**, N=30-110 different location measurements from 7-8 cross-sections. In panels **h** & **i**, N=38-74 villi were analyzed. *, p<0.05; **, p<0.01; ^#^, p<0.0001 using multiple *t-*test.

**Extended data Table 1.**
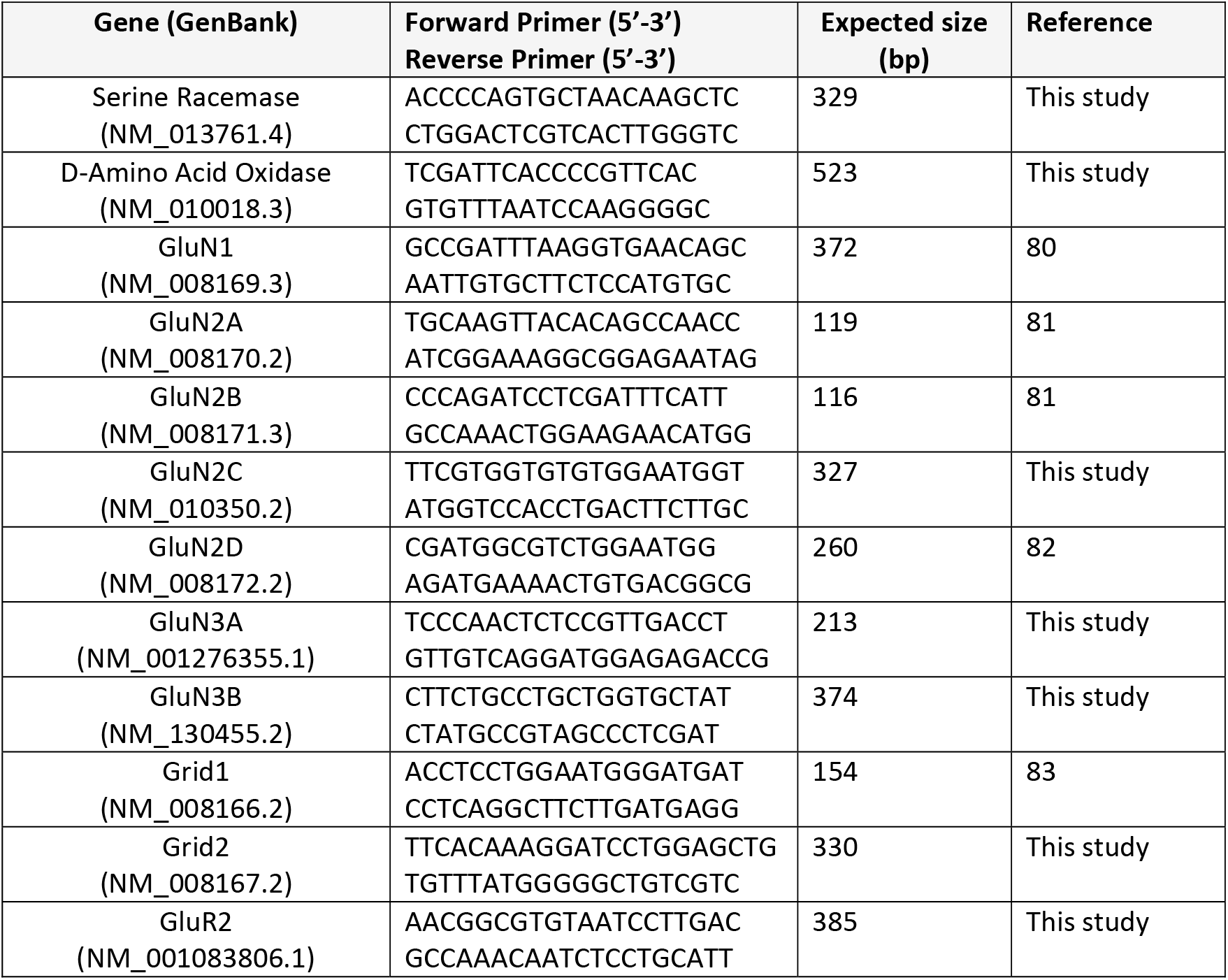
Sequences of primers and expected sizes of the amplicons.

**Extended data Table 2.** Antibody details.

| **Primary Antibody** | **Host species** | **Dilution /Use** | **Supplier** | **Catalog No** |
| --- | --- | --- | --- | --- |
| Monoclonal anti-Serine Racemase | Mouse | 1:400-/ WB<br>1:200 / IF | Santa Cruz Bio | sc-365217 |
| Polyclonal anti-Serine Racemase | Goat | 1:200 / IF | Santa Cruz Bio | sc-5752 |
| Polyclonal anti-D-Amino Acid Oxidase | Rabbit | 1:1,000/ WB | Jumpei Sasabe | Ref 41 |
| Polyclonal anti-D-Serine | Rabbit | 1:3K-1:5K/IF | GemacBio | AP041 |
| Polyclonal anti GluN1 C-ter | Goat | 1:500 / WB | Santa Cruz Bio | sc-1467 |
| Polyclonal GluN1 N-ter | Guinea Pig | 1:500 / WB | Alomone | AGP-046 |
| Polyclonal anti GluN2A | Rabbit | 1:1,000 / WB | Millipore | AB1555P |
| Polyclonal GluN2B | Rabbit | 1:750 / WB | Millipore | AB15362 |
| Polyclonal anti GluN2C/2D | Rabbit | 1:500 | Alomone | AGC-018 |
| Polyclonal anti-GluN3A | Rabbit | 1:500 / WB | Millipore | 07-356 |
| Polyclonal anti GluN3B | Rabbit | 1:500 / WB | Millipore | 07-351 |
| Polyclonal anti GluD1 | Rabbit | 1:10,000/WB | L Tricoire | Ref 84 |
| Polyclonal anti GluD2 C-ter | Rabbit | 1:500/WB | Frontier Inst Co Ltd | Rb-Af500 |
| Polyclonal anti GluD2 N-ter | Rabbit | 1:500 / WB | Frontier Inst Co Ltd | Rb-Af1460 |
| Polyclonal anti GluA2 | Rabbit | 1:400/WB | Alomone | AGC-005 |
| Polyclonal anti VGlut1 | Rabbit | 1:2,000/WB | Millipore | ABN1647 |
| Monoclonal anti β-Actin | Mouse | 1:10,000/WB | Sigma-Aldrich | A1978 |
| Monoclonal anti GAPDH | Rabbit (14C10) | 1:1,000 /WB | Cell signaling Tech | 2118 |
| Polyclonal anti-GFAP | Rabbit | 1:1,000/IF | DAKO | Z0334 |
| Monoclonal anti-S100β | Mouse | 1:1,000 / IF | Sigma-Aldrich | S2532 |
| Monoclonal anti-2H3 | Mouse | 1:5 / IF | DSHB | 2H3* |
| Monoclonal anti-Calbindin | Mouse | 1:100 / IF | Sigma-Aldrich | G9848 |
| Monoclonal anti-HUC/D | Mouse | 1:20 / IF | Molecular Probes | A21271 |
| Monoclonal anti-NeuN | Mouse | 1:1,000 / IF | AbCys | VMA377 |
| Monoclonal anti-NOS | Mouse | 1:500 / IF | Sigma-Aldrich | N2280 |
| **Secondary Antibody (HRP)** | **Host species** | **Dilution /Use** | **Supplier** | **Catalog No** |
| Polyclonal purified anti-rabbit | Goat | 1:4,000/WB | Abliance | BI2407 |
| Monoclonal purified anti-goat | Mouse | 1:4,000/WB | Abliance | B12403 |
| Polyclonal purified anti-mouse | Rabbit | 1:4,000/WB | Abliance | BI2413L |
| **Secondary Antibody (Fluo)** | **Host species** | **Dilution /Use** | **Supplier** | **Catalog No** |
| Alexa Fluor 546 anti-rabbit | Goat | 1:2,000/IF | Invitrogen | A11035 |
| Alexa Fluor 488 anti-mouse | Donkey | 1:1,000/IF | Invitrogen | A21202 |

1063 \*2H3 was deposited to the DSHB by Jessell, T.M. / Dodd, J. (DSHB Hybridoma Product 2H3).

