## Supplementary figures and images for "D-Serine agonism of GluN1-GluN3 NMDA receptors regulates the activity of enteric neurons and coordinates gut motility"

### Supplementary Figure 1

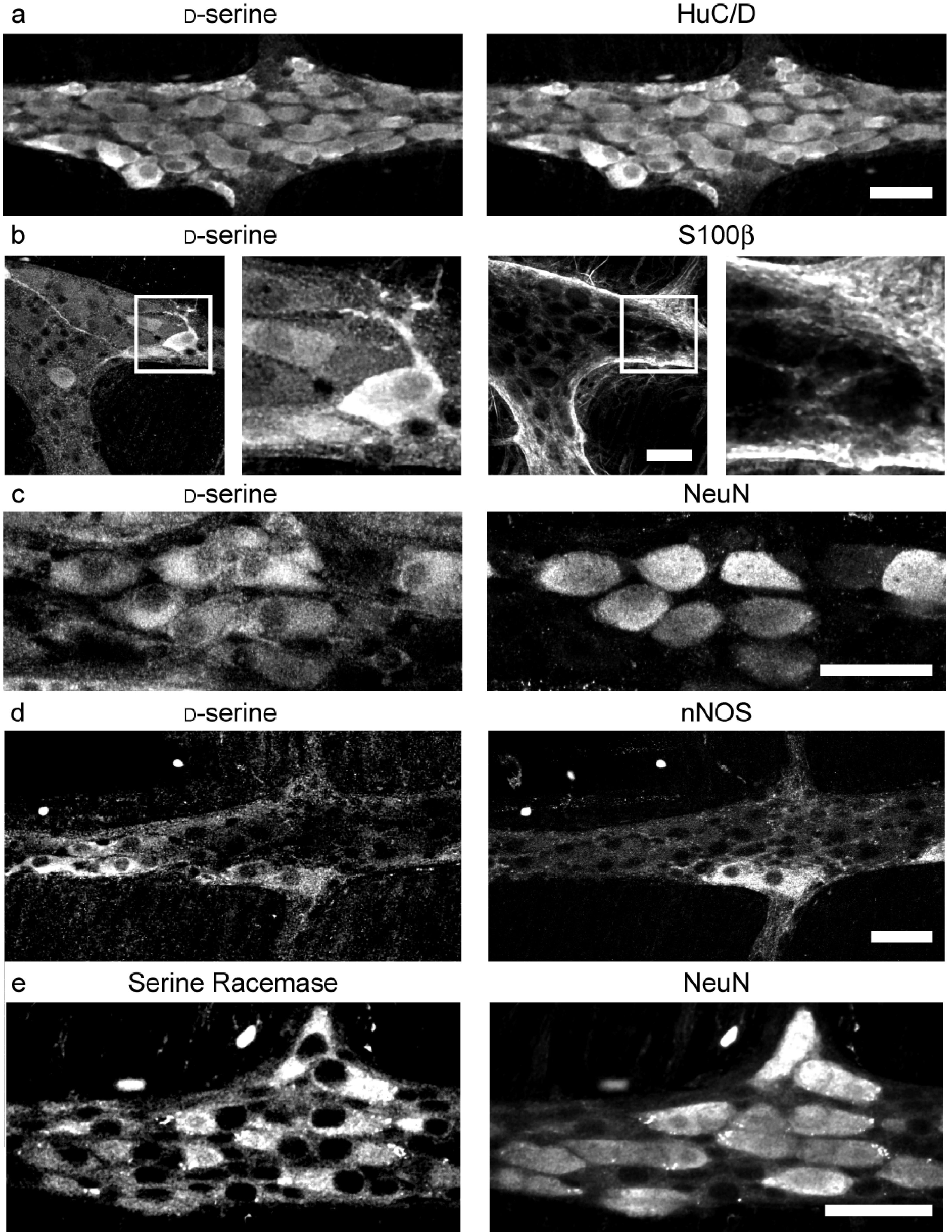

### Supplementary Figure 2

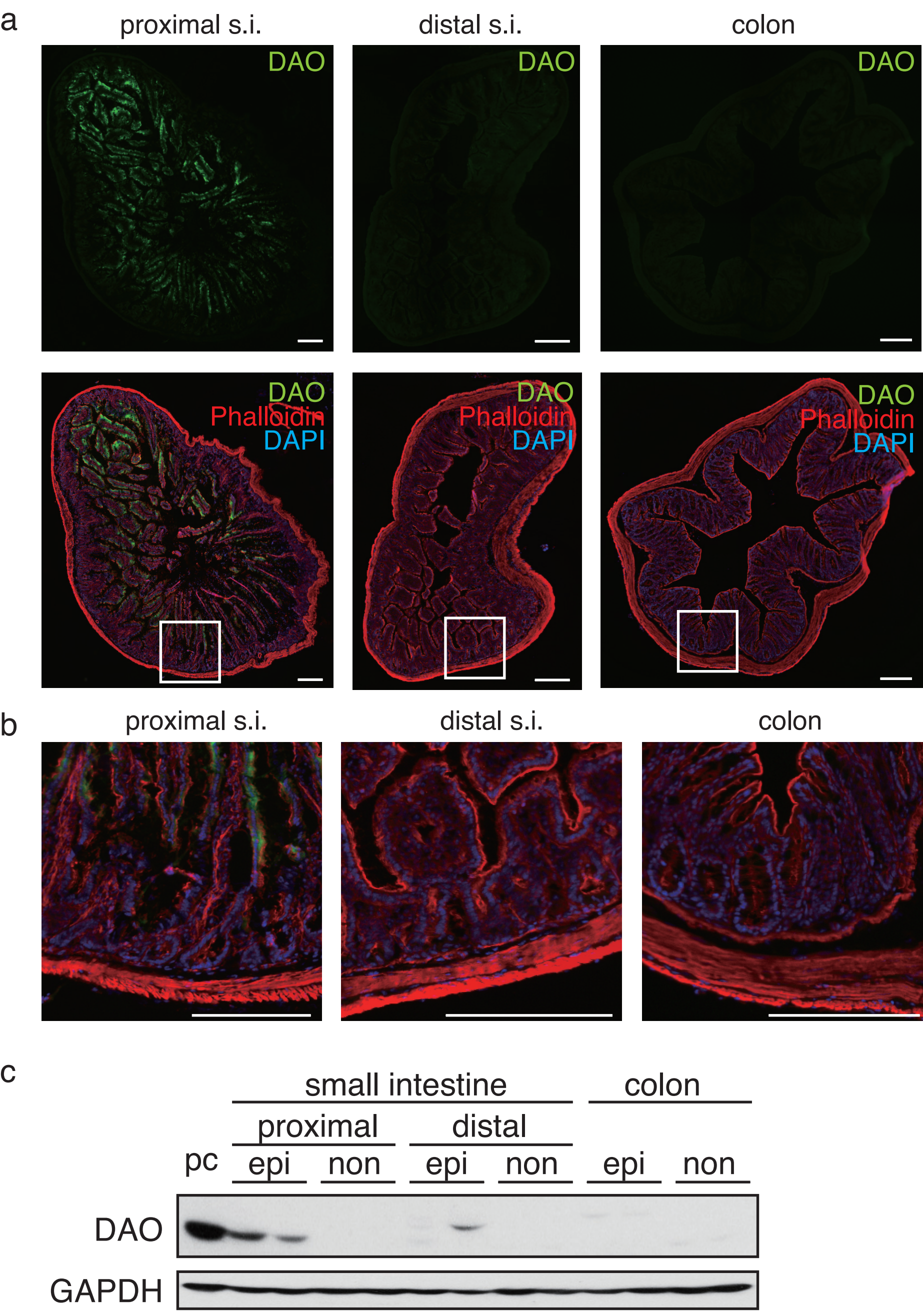

### Supplementary Figure 3

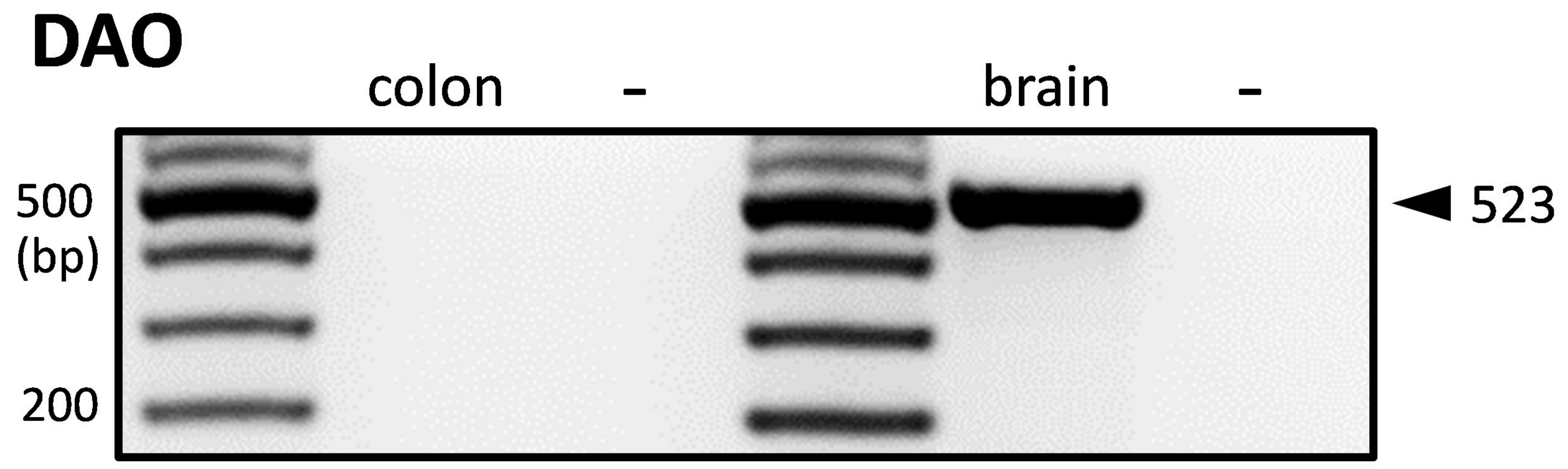

### Supplementary Figure 4

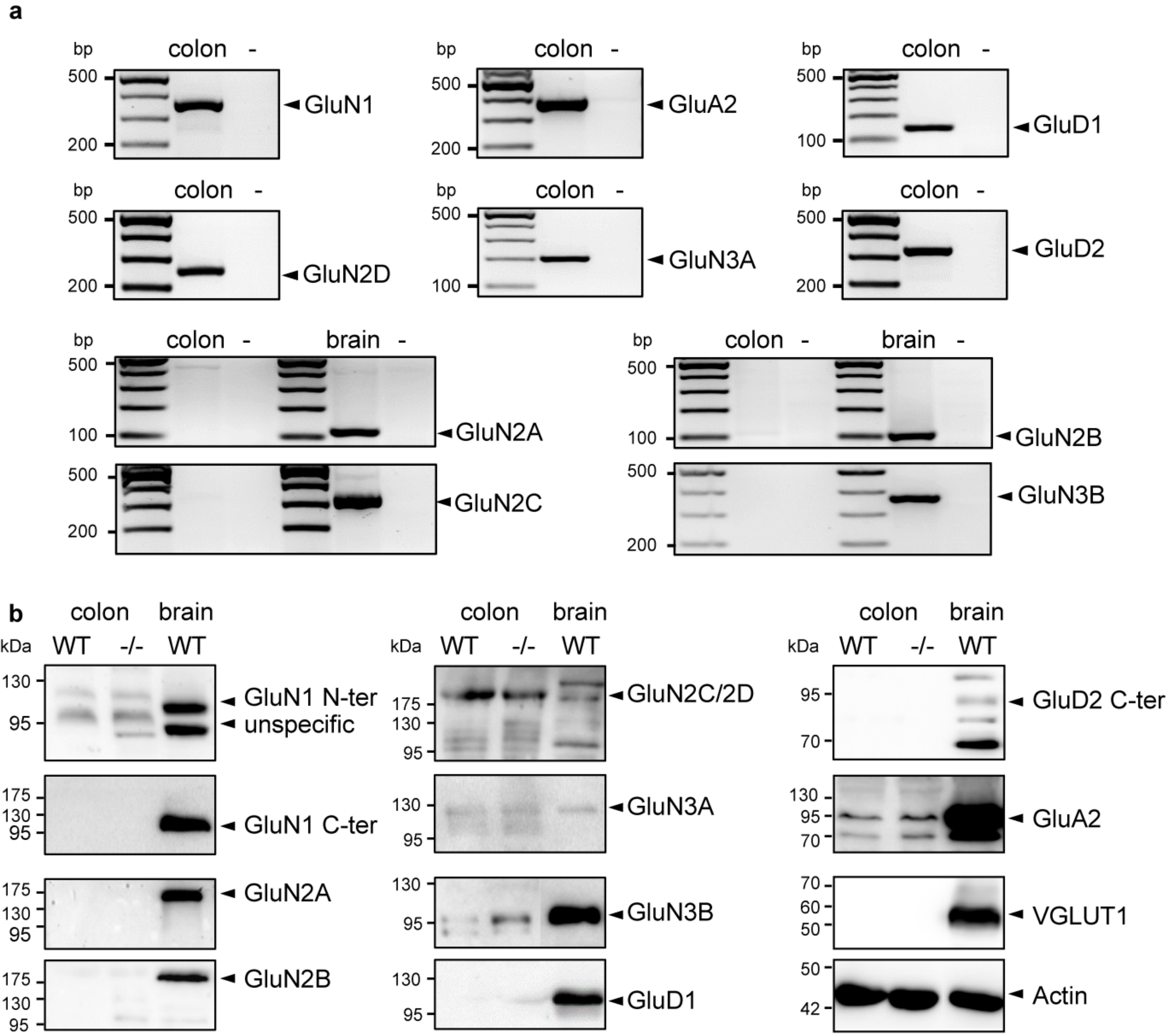

### Supplementary Figure 5

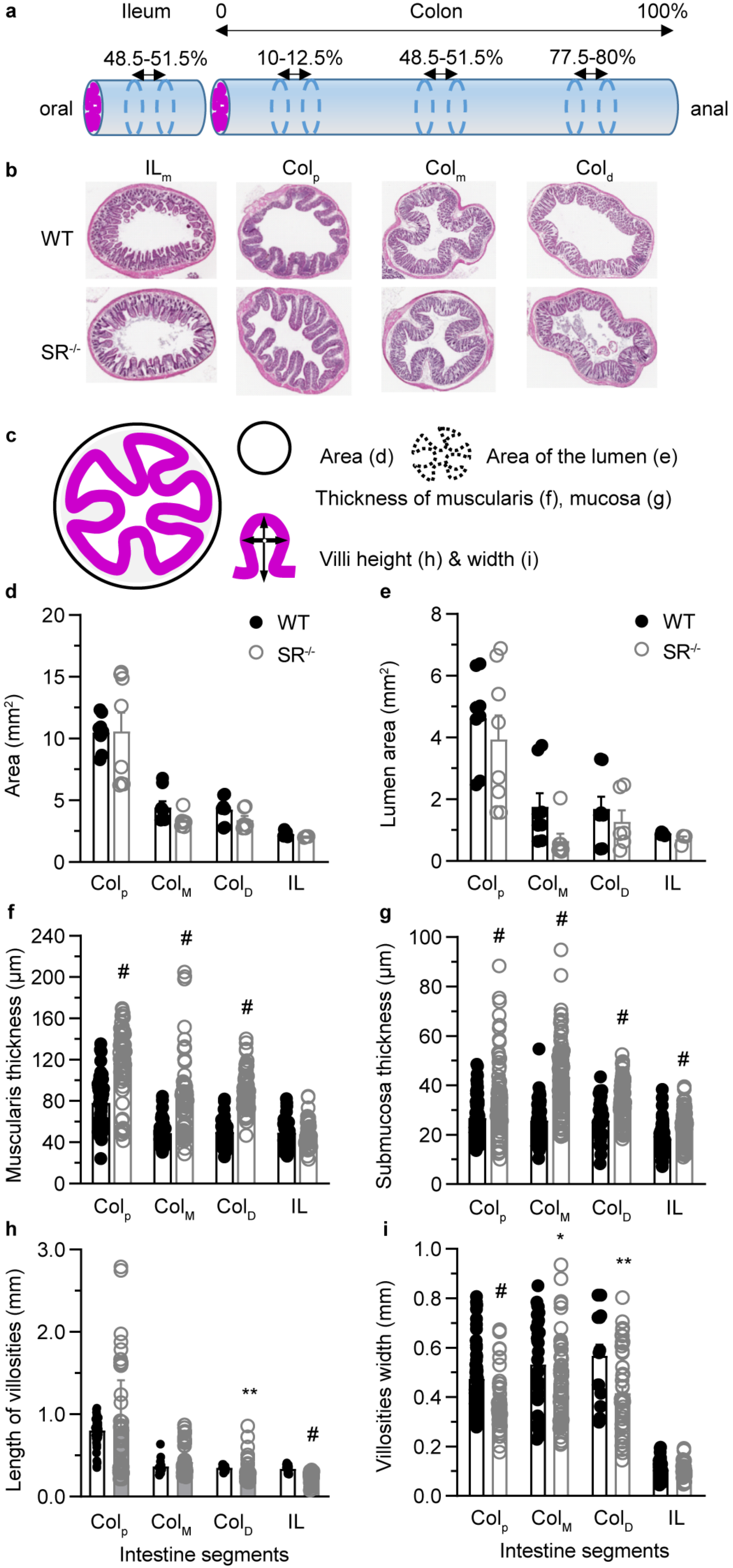
